# GGA2 and RAB13 promote activity-dependent ß1-integrin recycling

**DOI:** 10.1101/353086

**Authors:** Pranshu Sahgal, Jonna Alanko, Jaroslav Icha, Ilkka Paatero, Hellyeh Hamidi, Antti Arjonen, Mika Pietilä, Anne Rokka, Johanna Ivaska

## Abstract

β1-integrins mediate cell-matrix interactions and their trafficking is important in the dynamic regulation of cell adhesion, migration and malignant processes like cancer cell invasion. Here we employ an RNAi screen to characterize regulators of integrin traffic and identify the association of Golgi-localized gamma ear-containing Arf-binding protein 2 (GGA2) with β1-integrin and its role in recycling of the active but not inactive β1-integrin receptors. Silencing of GGA2 limits active β1-integrin levels in focal adhesions and decreases cancer cell migration and invasion congruent with its ability to regulate the dynamics of active integrins. Using the proximity-dependent biotin identification (BioID) method, we identify two RAB family small GTPases, RAB13 and RAB10, associating with GGA2 and β1-integrin. Functionally, RAB13 silencing triggers the intracellular accumulation of active β1-integrin, reduces integrin activity, in focal adhesions, and cell migration, similarly to GGA2 depletion, indicating that both facilitate active β1-integrin recycling the plasma membrane. Thus, GGA2 and RAB13 are important specificity determinants for integrin activity-dependent traffic.

## Introduction

Cancer cell migration is a multistep process that involves the spatiotemporal control of cell adhesion dynamics on multiple levels. Integrins are the main adhesion receptors for extracellular matrix (ECM) ligands. They consist of 18 α-and 8 β-subunits and can form 24 heterodimers for different extracellular matrix ligands (Hynes, 2002; Kawauchi, 2012). Integrin-mediated cell-ECM interactions are influenced by levels of integrin expression and integrin activity (i.e., affinity for ECM ligands) and by integrin endo/exosomal trafficking on the plasma membrane. Integrin activity in adherent cell types, like carcinoma cells and fibroblasts, is most likely defined by a continuum of receptor conformations where the fully inactive bent receptor switches in response to integrin activating proteins, such as talin and kindlin, into the fully active, extended and ligand-bound receptor (Ye et al., 2014; Calderwood et al., 2013). Deregulated integrin activity is implicated in multiple pathological conditions (Winograd-Katz et al., 2014; Pozzi and Zent, 2013; Hegde and Raghavan, 2013). In cancer cells, increased integrin activity, triggered either by increased expression of integrin activating proteins (Calderwood et al., 2013) or reduced expression of inactivating proteins (Liu et al., 2015; Rantala et al., 2011b), has been linked to increased migration and invasion (Jin et al., 2015; Felding-Habermann et al., 2001).

Integrins undergo constant endo/exocytosis from the cell membrane and this is important to provide directionality to the cell and to assist in the formation and release of cell-substrate contacts for proper cell movement (Moreno-Layseca et al., 2019; Kawauchi, 2012; Fletcher and Rappoport, 2010). Thus, the balance between internalized and cell-surface integrins within cells plays a critical role in cell migration. Altered integrin trafficking, especially in the case of β1-integrins, is linked to invasive processes (Bridgewater et al., 2012; De Franceschi et al., 2015; Parachoniak and Park, 2012). Furthermore, recent studies have reported the presence of active integrins inside endosomes (Alanko et al., 2015; Rainero and Norman, 2015) and a link between directed integrin traffic, from talin and FAK-positive endosomes to the plasma membrane, and the reassembly of focal adhesions at the leading edge of migrating cells (Nader et al., 2016). Previous work, from us, has shown that the net endocytosis rate of active β1-integrins is higher than inactive β1-integrins (Arjonen et al., 2012). Active β1-integrins recycle predominantly through RAB11-dependent long-loop pathways whereas inactive β1-integrins recycle through RAB4 and actin-dependent short-loop recycling pathways or undergo retrograde traffic through the trans-Golgi network (Powelka et al., 2004; Arjonen et al., 2012; Campbell and Humphries, 2011). However, apart from some more specific conditions such as RAB25-and CLIC3-expressing carcinomas (Dozynkiewicz et al., 2012), the mechanistic details of the trafficking pathways that differentially regulate integrin dynamics based on receptor activity remain poorly understood. Furthermore, very little is known about how different endosomal adaptor proteins contribute to integrin traffic and cell migration.

The GGA (Golgi-localized gamma ear-containing Arf-binding protein) family of adaptors (GGA1, GGA2, GGA3) was first described in the year 2000 and originally identified as Arf effectors, interacting with clathrin-coated vesicles and localizing to the trans-Golgi network and endosomes (Boman et al., 2000; Dell’Angelica et al., 2000; Puertollano et al., 2001; Puertollano and Bonifacino, 2004). Even though the mammalian GGAs are thought to be at least partially redundant (Hirst et al., 2000; Dell’Angelica et al., 2000; Govero et al., 2012), several studies have indicated specific non-overlapping and even opposite functions for GGA-family members. In mice, genetic deletion of GGA1 or GGA3 alone is well tolerated whereas the loss of GGA2 results in embryonic or neonatal lethality (Govero et al., 2012), indicating important and specific biological functions for GGA2. Moreover, GGA1 and GGA3, but not GGA2, recognize ubiquitin (Puertollano and Bonifacino, 2004; Scott et al., 2004; Shiba et al., 2004) and have been implicated in the degradation of endocytosed EGFR. While GGA3 or GGA1 depletion promotes EGFR expression, GGA2 depletion, in contrast, drastically reduces EGFR levels (Uemura et al., 2018). This could be linked to the fact that GGA2 interacts with and stabilizes activated EGFR (O’Farrell et al., 2018). In addition, GGA1 and GGA3 are specifically required for endosomal recycling of transferrin (GGA1) and MET receptor tyrosine kinase (GGA3) and in facilitating stability of collagen-binding integrin α2β1 (GGA3) by diverting it away from lysosomal degradation (Zhao and Keen, 2008; Parachoniak et al., 2011; Ratcliffe et al., 2016). In contrast, GGA2 has not been implicated in integrin traffic.

Here, we have investigated the role of the GGA adaptor family in regulating β1-integrin traffic in breast cancer cells and identify a previously undescribed role for GGA2 and RAB13 in driving recycling of active β1-integrins and promoting cell motility and invasion.

## Results

### GGA2 regulates β1-integrin traffic

The coordinated traffic of integrins through the endolysosomal network must be tightly regulated to enable efficient cell motility. Unlike many other cell-surface receptors, integrins have a remarkably long half-life suggesting that following endocytosis the majority of integrins are efficiently recycled back to the plasma membrane. We recently identified a critical role for GGA3 in diverting the collagen receptor α2β1-integrin away from lysosomal degradation to support HeLa cell migration on collagen (Ratcliffe et al., 2016). This prompted us to investigate whether other GGAs or their known interactors (based on a BioGRID server search) regulate integrin traffic. We conducted a β1-integrin endocytosis screen employing a quenching-based β1-integrin antibody internalization assay (Arjonen et al., 2012) adapted to the cell spot microarray RNAi screening platform (Rantala et al., 2011a). Focusing on GGAs and their interactors, we consistently observed a significant increase in intracellular β1-integrin levels, after 30 minutes of endocytosis, upon GGA2 silencing with two independent siRNAs in MDA-MB-231 cells (Fig. S1A). To confirm the role of GGA2 in β1-integrin dynamics, we performed biotin-IP-based β1-integrin endocytosis assays in GGA2-silenced (GGA2 siRNA#1 is distinct from the RNAi oligos used in the screen) MDA-MB-231 cells. In line with the RNAi screen, depletion of GGA2 significantly increased the level of intracellular β1-integrin at 10 and 20 minute time points without influencing total β1-integrin levels (Fig. 1A,B; S1B). Conversely, overexpression of GFP-GGA2 significantly reduced β1-integrin uptake (Fig. 1C,D). In contrast to GGA1 depletion (Zhao and Keen, 2008), silencing of GGA2 did not influence internalization of the transferrin receptor, suggesting a specific role for GGA2 in integrin receptor traffic (Fig. S1C,D).

**Figure 1.**
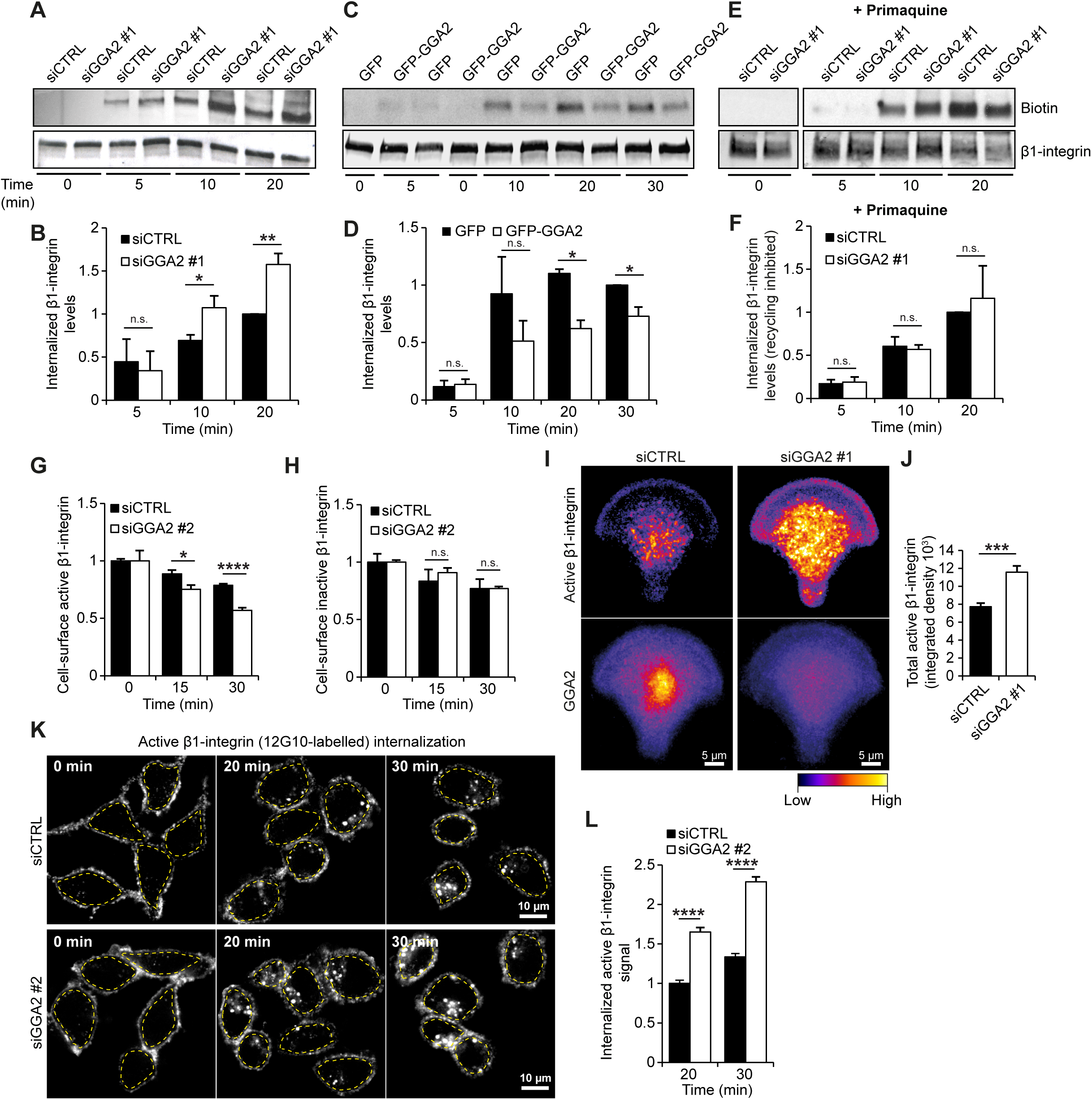
GGA2 regulates β1-integrin receptor traffic. **A-B)** Internalization of biotinylated cell-surface β1-integrin in MDA-MB-231 cells treated with either control siRNA (siCTRL) or GGA2 siRNA oligo #1 (siGGA2 #1). Shown is a representative western blot (A) and quantification of biotinylated β1-integrin relative to total β1-integrin levels and normalized to siCTRL 20 min time point (B). **C-D)** Internalization of biotinylated cell-surface β1-integrin in MDA-MB-231 cells expressing GFP or GFP-GGA2. Shown is a representative western blot (C) and quantification of biotinylated β1-integrin relative to total β1-integrin and normalized to GFP at 30 min time point (D). **E-F)** Biotinylation-based integrin internalization assays performed, analyzed and normalized as in (A,B) but in the presence of the recycling inhibitor primaquine (100 µM). **G-H)** Flow cytometry analysis of cell-surface labelled active (12G10 antibody) (G) or inactive (MAB13 antibody) (H) β1-integrin cell-surface levels in MDA-MB-231 cells treated with siCTRL or siGGA2 #2 (GGA2 siRNA oligo #2), at the indicated internalization periods, normalized to siCTRL at 0 min time point. **I-J)** Representative heatmaps (I) of the mean fluorescence intensity of active β1-integrin (9EG7 antibody) and GGA2 in siCTRL or siGGA2#1 MDA-MB-231 cells plated on fibronectin-coated crossbow-shaped micropatterns. Total levels of active β1-integrin in each condition were also quantified (J; data are mean ± SEM; n = 38 cells in each condition). **K)** Representative confocal microscopy images of active β1-integrin (12G10 antibody) at the indicated time points in siCTRL or siGGA2 #2 MDA-MB-231 cells. **L)** Quantification of intracellular active β1-integrin levels. The intracellular fluorescence signal was quantified within the area defined by a dashed yellow outline (in K), which excludes any signal from the plasma membrane. The measured signal for each time point was then normalized to the background fluorescence at 0 min from the respective experimental condition, i.e. the signals from siCTRL time points were normalized to 0 min siCTRL and signals from siGGA2 time points were normalized to 0 min siGGA2 (data are mean ± SEM; n = 100-120 cells per condition). A-H; Data are mean ± SEM of n = 3 independent experiments. Statistical analysis, Student’s unpaired *t*-test (n.s.= not significant; *p < 0.05; **p < 0.005; ***p < 0.0005; ****p < 0.00001).

### GGA2 promotes specifically the recycling of active β1-integrin

Many studies have indicated that β1-integrins recycle to the plasma membrane with a turnover rate of 10-15 minutes (Argenzio et al., 2014; Diggins et al., 2018; Dozynkiewicz et al., 2012). The slow accumulation of β1-integrin inside GGA2-silenced cells led us to hypothesise that receptor recycling, rather than endocytosis, is affected by the loss of GGA2 expression. Indeed, the recycling inhibitor primaquine fully abolished the observed difference in β1-integrin uptake between control-and GGA2-silenced cells (Fig. 1E,F), indicating that GGA2 silencing increases intracellular β1-integrin levels by inhibiting recycling rather than stimulating endocytosis.

To confirm the specificity of these findings and the implications for active and inactive pools of β1-integrins, we silenced GGA2 with a fourth independent siRNA (GGA2 siRNA#2 is distinct from the RNAi oligos used in the screen) and analyzed the levels of cell-surface labelled active and inactive β1-integrins, following internalization, using flow cytometry (Fig. 1G,H). Interestingly, loss of GGA2 was accompanied by a significant reduction in the cell-surface levels of the active, but not inactive, β1 integrin receptor after endocytosis was triggered (at 15 and 30 min time points) (Fig. 1G,H), suggesting selective intracellular accumulation of the active receptor in the absence of GGA2. Further immunofluorescence analyses of control-and GGA2-silenced cells plated on micropatterns (Théry, 2010) supported the results obtained with the biotin-and flow cytometry-based integrin trafficking assays, and demonstrated a significant increase in the steady-state pool of active β1-integrin in endosome-like structures in the cytoplasm upon GGA2 silencing (Fig. 1I,J). Immunofluorescence of internalized cell-surface-labelled active β1-integrin, following 20 or 30 minute antibody pulse, corroborated a significant increase in the intracellular accumulation of active β1-integrin in endosome-like structures in GGA2-silenced cells (Fig. 1K,L). This was specifically due to loss of GGA2 and not off-target effects as re-expression of GGA2, in cells silenced with a GGA2 3’ UTR targeting oligo, returned integrin endocytosis to control levels (Fig. 2A,B). These puncta of internalized active-β1 were significantly reduced by overexpression of GFP-GGA2 (Fig. 2C,D) and they partially overlapped with GFP-GGA2 (Fig. 2C). In contrast, GFP-GGA2 expression had no significant effect on internalization of cell-surface-labelled inactive-β1 integrin (Fig. S1E, F). Inhibition of receptor recycling with primaquine abolished the difference in active β1-integrin internalization between GFP and GFP-GGA2 expressing cells (Fig. 2E,F), further validating a role for GGA2 in recycling of active β1-integrin in these cells. Interestingly, GFP-pulldowns of GFP-GGA2, but not GFP alone, co-precipitated endogenous β1-integrin (Fig. 2G). GFP-GGA2 association with β1-integrin was further validated by repeating the pulldown in another breast cancer cell line (MCF10DCIS.com) and confirming that GFP-GGA2 pulls down its known interactors, clathrin and GGA3, in addition to β1-integrin (Fig. 2H). These data suggest that GGA2 and β1-integrin may exist in the same molecular complex within cells and in this way GGA2 may facilitate integrin trafficking.

**Figure 2.**
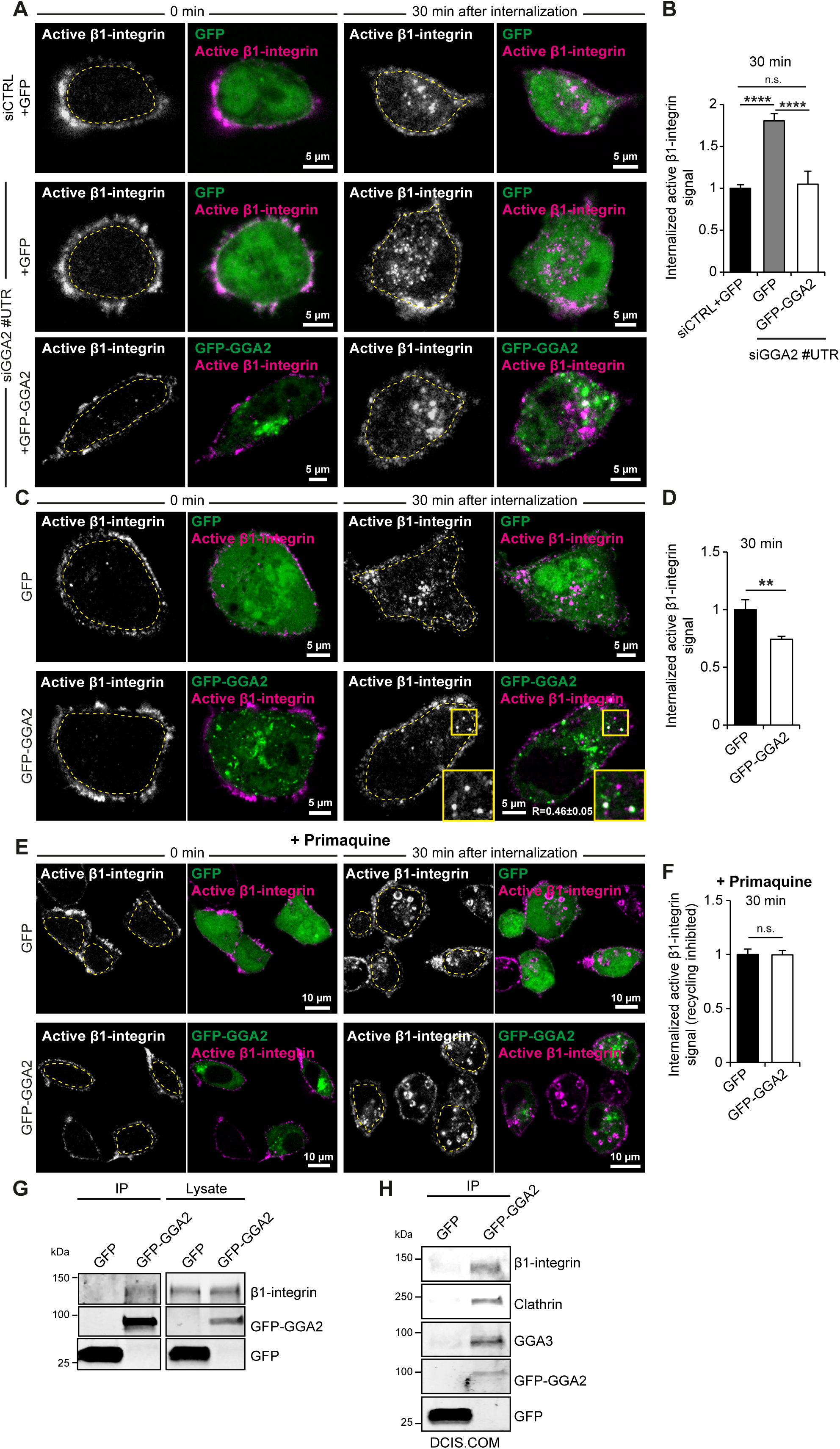
GGA2 associates with β1-integrins and promotes recycling of active β1-integrins. **A-B)** Representative confocal microscopy images (A) and quantification (B, performed as in Fig. 1L) of internalized active β1-integrin (cell-surface labelled with 12G10 antibody) at the indicated time points in MDA-MB-231 cells transfected with either siCTRL or siRNA targeting the 3’ untranslated region (UTR) of GGA2 and re-expressing GFP or GFP-GGA2 (data are mean ± S.E.M; n = 55-95 cells per condition). **C-D)** Representative confocal microscopy images (C) and quantification (D, performed as in Fig. 1L) of internalized active β1-integrin (12G10 antibody) at the indicated time points in MDA-MB-231 cells expressing GFP or GFP-GGA2. Overlap between GFP-GGA2 and active β1-integrin is quantified (R, Pearson correlation coefficient between fluorescence signals in the two channels, ± SD; n = 109 cells). Intracellular active β1-integrin levels (30 min) were normalized to GFP MDA-MB-231 cells at 0 min time point (data are mean ± S.E.M; n = 70-110 cells per condition, D). **E-F)** Representative images (E) and quantification (F, performed as in Fig. 1L) of internalized active β1-integrin (12G10 antibody) at the indicated time points in MDA-MB-231 cells expressing GFP or GFP-GGA2 and treated with 100 µM primaquine (data are mean ± SEM; n = 70-80 cells per condition). **G)** Representative western blot analysis of GFP-Trap pulldowns performed in GFP or GFP-GGA2 expressing MDA-MB-231 cells (n = 3 independent experiments). **H)** Representative western blot analysis of GFP-Trap pulldowns performed in MCF10ADCIS.COM breast cancer cells transiently transfected with GFP or GFP-GGA2. GGA2-interacting proteins GGA3 and clathrin are shown as positive controls. Yellow insets show magnified regions of interest (ROI). Statistical analysis, Student’s unpaired *t*-test (n.s. = not significant; *p < 0.05; **p < 0.005; ***p < 0.0005; ****p < 0.00001).

### GGA2 localizes to early and recycling endosomes

GGA2 has been studied predominantly in yeast and its subcellular localization in cancer cells is less well characterized. We performed immunofluorescence staining of endogenous GGA2 and markers of different endomembrane compartments. GGA2 overlapped with early endosomal marker RAB5 and with recycling endosomes positive for RAB4 or RAB11 (Fig. 3A 1-3.). In addition, partial overlap was detected with early endosomal marker EEA1 (Fig. 3A 4.) and with VPS35, a component of the retromer complex (Fig. 3A 5.). GGA2 did not overlap with late endosome/lysosome markers RAB7 and LAMP1 (LAMP1 was stained in GFP-GGA2-expressing cells, as antibodies against LAMP1 and GGA2 are raised in the same species) (Fig. 3A 6-7). Similar subcellular localization was also observed for GFP/RFP-tagged GGA2 (Fig. S2A). Thus, in MDA-MB-231 cells, GGA2 in addition to its established localization to TGN46-labelled trans-Golgi network, RAB6-positive Golgi vesicles and clathrin-coated vesicles (Fig. S2B,C), is also found in early and recycling endosomes (Fig. S2A,D and Fig. 3A) (Zhu et al., 2001; Hirst et al., 2000). Moreover, and in support of a role in integrin trafficking, we found partial overlap between GGA2 and β1-integrin in intracellular vesicles (β1-integrin was stained in GFP-GGA2-expressing cells, as antibodies against β1-integrin and GGA2 are raised in the same species) (Fig. 3B). Thus, GGA2 partially overlaps with β1-integrin, localizes to early and recycling endosomes.

**Figure 3.**
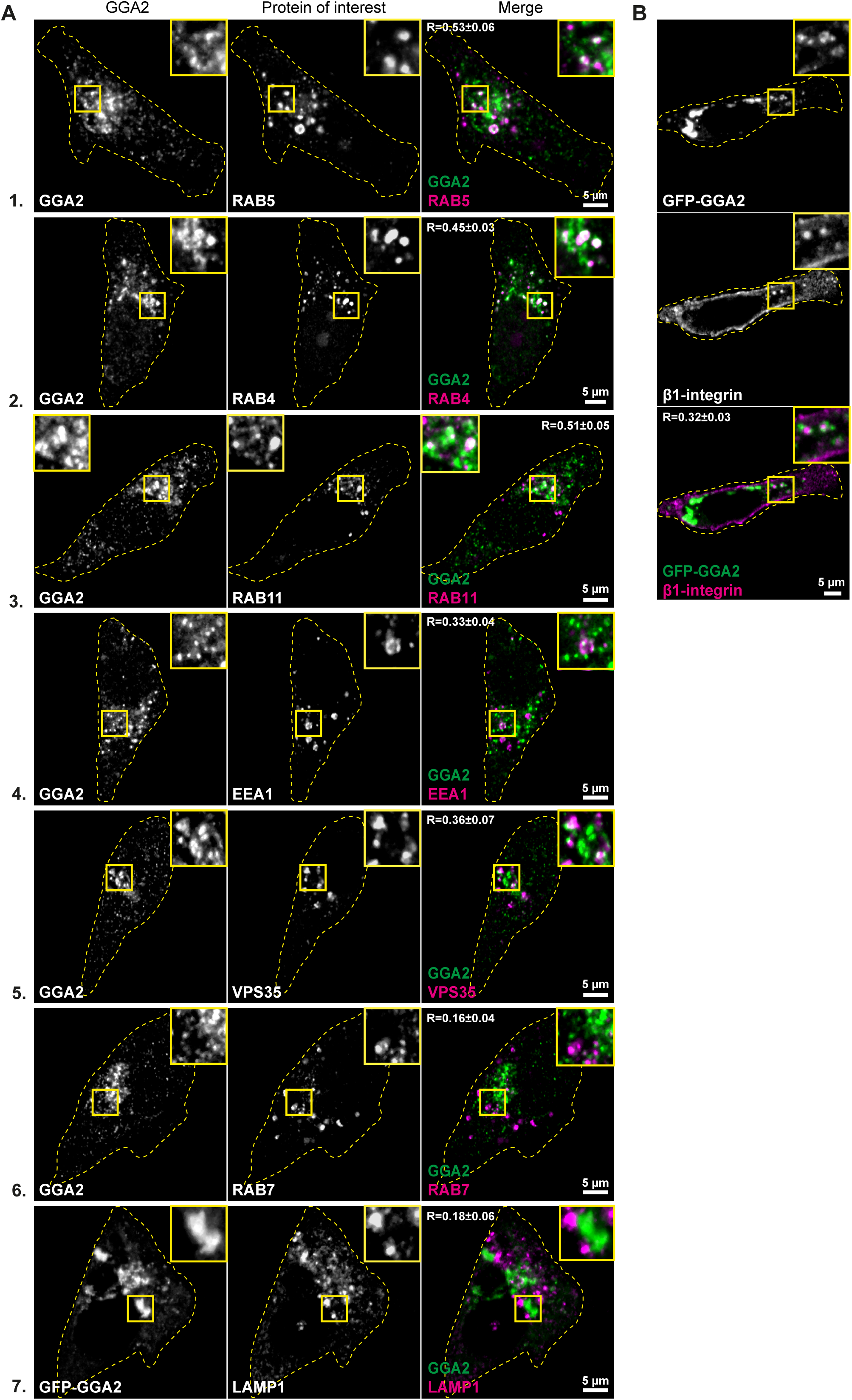
Cellular localization of GGA2. **A)** Representative confocal microscopy images of MDA-MB-231 cells co-stained with antibodies against the indicated vesicular compartment proteins (proteins of interest, magenta) and an antibody against endogenous GGA2 (1-6). LAMP1 staining was analyzed in cells expressing GFP-GGA2 (7). **B)** Representative confocal microscopy images of MDA-MB-231 cells expressing GFP-GGA2 and stained for endogenous β1-integrin (P5D2; total β1-integrin). For (A) and (B) overlap of signals are quantified (R ± SD; n = 80-200 cells per condition). Yellow dashed lines delineate cell outlines. Yellow insets show magnified ROI.

### GGA2 is a positive regulator of cell migration and cancer cell invasion

β1-integrins and their endo/exocytic traffic are key regulators of cell-ECM interactions and cell motility. To investigate whether GGA2 regulates focal adhesions, we plated control-and GGA2-silenced cells on a mixture of collagen and fibronectin and analysed the number and size of paxillin-positive focal adhesions. While loss of GGA2 had no significant effect on focal adhesion number (Fig. 4A), it significantly reduced active β1-integrin levels in paxillin-positive adhesions 30 minutes after plating and the lower active β1-integrin levels were maintained up to 2 hours post plating (Fig. 4B, C). Recycling of β1-integrins is essential for cell migration and for effective cancer cell invasion (Paul et al., 2015a; Caswell et al., 2008). However, most integrin trafficking pathways described thus far do not distinguish between active and inactive integrin receptors (Paul et al., 2015b; Moreno-Layseca et al., 2019). Therefore, we wanted to determine whether the specific inhibition of active β1-integrin recycling could influence cancer cell motility and invasion. Time-lapse microscopy revealed significantly slower MDA-MB-231 cell migration following GGA2 silencing (Fig. 4D). Furthermore, GGA2 depletion reduced cancer cell invasion in three-dimensional collagen (Fig. 4E). These effects were specifically due to reduced cell motility and altered adhesions since GGA2 silencing did not influence cell proliferation in 2D or in suspension (Fig. S3A,B). Together, these results show that interfering with the GGA2-dependent delivery of active β1-integrins to the plasma membrane leads to reduced cell motility and impaired cancer cell invasion in vitro.

**Figure 4.**
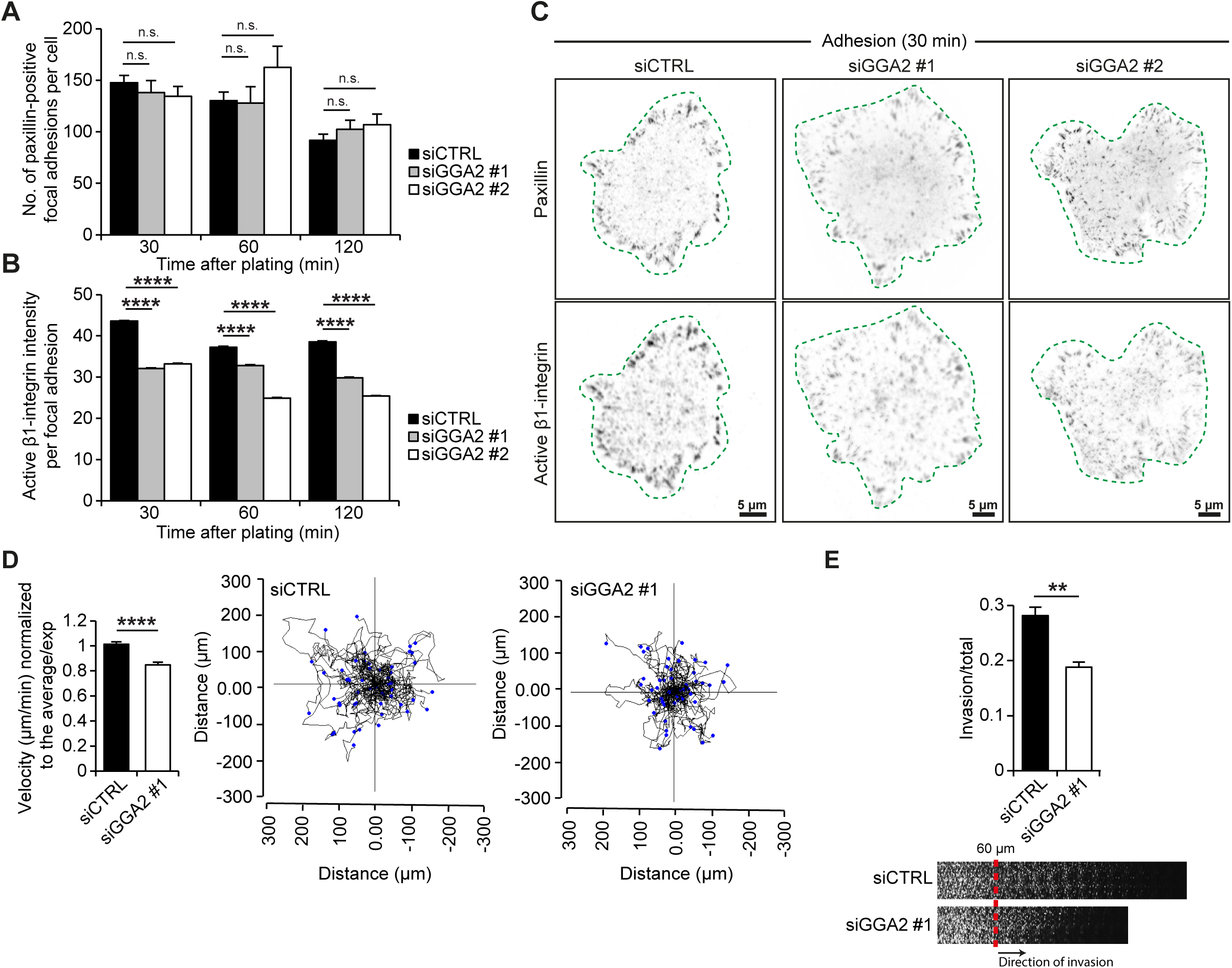
Loss of GGA2 reduces active β1-integrin levels in focal adhesions, cell migration and cancer cell invasion. **A-C)** siCTRL, siGGA2 #1 or siGGA2 #2 MDA-MB-231 cells were plated on collagen and fibronectin for the indicated time points, fixed and stained for paxillin (to detect focal adhesions) and active β1-integrin (12G10). Shown are quantifications of focal adhesion number (A) and 12G10 mean fluorescence intensity in focal adhesions (B) with representative confocal microscopy images (bottom plane) (C) (data are mean ± SEM; n = 7,700-15,700 focal adhesions from 80-110 cells per condition). Green dashed lines delineate cell outlines. **D)** Quantification of cell velocity in siCTRL or siGGA2 #1 MDA-MB-231 cells migrating randomly on fibronectin-coated dishes (data are mean ± SEM; n = 150-180 cells for each condition). Representative migration tracks are shown in black for 50 cells per condition. End-points are represented by blue dots. **E)** Inverted invasion assay showing reduced MDA-MB-231 cell invasion following GGA2 silencing. Invasion in collagen plugs supplemented with fibronectin was visualized using confocal microscope by imaging serial optical sections at 15 μm intervals. Individual confocal images are shown in sequence with increasing penetrance from left to right. The invasion was quantified using ImageJ by measuring the fluorescence intensity of cells invading further than 60 μm relative to the fluorescence intensity of all cells in the plug (data are mean ± SEM, n = 3 independent experiments). A-B, Student’s unpaired *t*-test; D-E, non-parametric Mann-Whitney test (n.s.= not significant; *p < 0.05; **p < 0.005; ***p < 0.0005; ****p < 0.00001).

### Proximity biotinylation identifies new GGA2-associated trafficking proteins

All GGA-family members share the VHS domain, which binds to cytosolic acidic di-leucine (AC-LL) motif-containing cargo proteins. Interactions at this site have been demonstrated to regulate trafficking of cargo such as the mannose-6-phosphate receptor and sortilin (Takatsu et al., 2001; Nielsen et al., 2001). β1-integrins lack the AC-LL motif, suggesting that alternative mechanisms are involved in the GGA2-dependent traffic of active β1-integrins. To investigate this further, we employed in-situ labelling of GGA2-proximal proteins in intact cells by fusing GGA2 with BirA (BioID) (Fig. S3C) and myc-tag to enable visualization of the protein (Roux et al., 2012). Importantly, Myc-BioID-GGA2 localized similarly as GFP-GGA2 to a perinuclear compartment (Golgi) and to a pool of endosomes and efficiently catalyzed in-situ biotinylation of these compartments in biotin-treated cells (Fig. S3D). In contrast, Myc-BioID alone did not overlap with GFP-GGA2 in cells and displayed uniform, non-compartmentalized biotinylation of the cytosol (Fig. S3D). The specificity and efficiency of this approach was further validated by isolating biotinylated proteins from Myc-BioID-or Myc-BioID-GGA2-expressing cells. Known GGA2 interacting proteins including clathrin, GGA1 and GGA3 (Zhu et al., 2001; Ghosh et al., 2003) were detected in streptavidin pulldowns only in Myc-BioID-GGA2-expressing cells, indicating functionality and specificity of the in-situ biotinylation approach (Fig. S3E).

To identify putative GGA2 binding proteins, four independent BioID experiments were carried out in MDA-MB-231 cells expressing Myc-BirA or Myc-BirA-GGA2 (Fig. S3D). Biotinylated proteins were detected by a semi-quantitative spectral counting mass spectrometry method. Proteins with higher number of peptide spectrum matches (PSM) in Myc-BirA-GGA2 than Myc-BirA in at least three experiments were considered as putative GGA2 binders (Table S1). The nine identified proteins included some known GGA2 interactors (ARF6 and IGFR2; Fig. 5A) as well as new putative GGA2-proximal proteins representing potentially new players in integrin traffic. As expected, gene ontology pathway analysis of the putative GGA2 interactors indicated significant enrichment of the trans-Golgi network. In addition, “recycling endosome membrane pathway” was significantly enriched based on three proteins (Fig. 5B); ARF6 is a known regulator of integrin recycling (Powelka et al., 2004; Fang et al., 2010; Schweitzer et al., 2011; Morgan et al., 2013), which has also been implicated in integrin endocytosis (Dunphy et al., 2006; Chen et al., 2014; Sakurai et al., 2010), whereas, RAB10 and RAB13, to the best of our knowledge, have not been previously linked to β1-integrin traffic (RAB13 has been implicated in Mst1 kinase-induced delivery of lymphocyte function-associated antigen-1, an immune-cell-specific integrin, to the cell surface) (Nishikimi et al., 2014). In addition, SEC23B, which was identified using BioID but did not quite reach the same confidence level as RAB10 and RAB13, was included for further investigation since in the interaction network, RAB13 and RAB10 formed a unique node (dotted box) with SEC23B as a common interactor (Fig. 5A). SEC23B is a component of the COPII responsible for creating small membrane vesicles that originate from the endoplasmic reticulum (ER) (Barlowe et al., 1994; Gorelick and Shugrue, 2001) and it associates with the RAB10 effector SEC16A in COPII-coated vesicles to regulate insulin-stimulated GLUT4 trafficking in adipocytes (Bruno et al., 2016). To validate the BioID results, we probed GFP-GGA2 GFP-Trap pulldowns for endogenous RAB10, RAB13 and SEC23B and found that these proteins were expressed in MDA-MB-231 cells and associated with GFP-GGA2 (Fig. 5C,D, Fig. S3F). Furthermore, immunofluorescence analyses demonstrated that endogenous RAB13 and GGA2, as well as GFP-RAB13 and GGA2, localize to overlapping perinuclear and plasma membrane-proximal vesicles in MDA-MB-231 cells (Fig. 5E, S3G). This is congruent with the established role of RAB13 function in exocytic membrane traffic from the trans-Golgi network and recycling endosomes to the cell surface in polarized epithelial cells and cancer cells (Nokes et al., 2008). While we were unable to identify antibodies for staining endogenous RAB10 or SEC23B, we observed overlap between GFP-RAB10 and GFP-SEC23B with endogenous GGA2 on cytoplasmic endomembranes (Fig. 5F, S3H). However, unlike RAB13, RAB10 and SEC23B were not detected on the plasma membrane.

**Figure 5.**
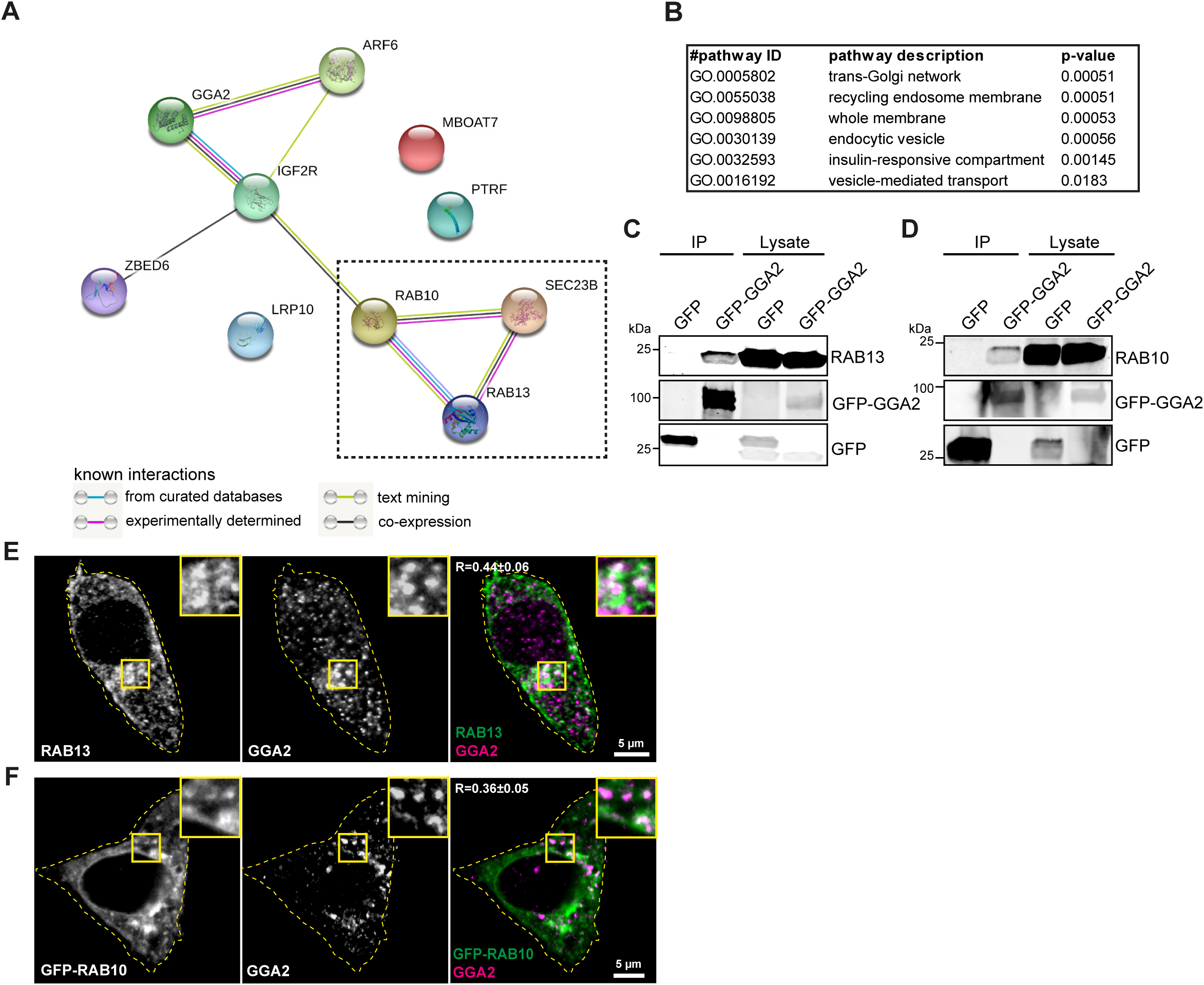
Proximity biotinylation identifies new GGA2-associated trafficking proteins. **A)** Interaction network of putative GGA2 proximal proteins identified with BioID created by STRING server using medium stringency (confidence 0.200). The node of interest is indicated by a dotted outline. **B)** Gene ontology analysis created by STRING server of the proteins depicted in (A). p-values indicate the false discovery rate provided by STRING. **C-D)** Representative western blot of GFP-Trap pulldowns in MDA-MB-231 cells transiently transfected with GFP or GFP-GGA2 and analyzed for GFP and endogenous RAB13 (C) or RAB10 (D) (n = 3 independent experiments). **E-F)** Representative confocal microscopy images of MDA-MB-231 cells stained for endogenous RAB13 and GGA2 (E) or transfected with GFP-RAB10 and stained for GGA2. Overlap between the indicated proteins is quantified (R ± SD; n = 80-200 cells per condition). Yellow insets show ROI.

### RAB13, RAB10 and SEC23B associate with β1-integrins

Next, we investigated the ability of GGA2 proximal proteins, RAB10, RAB13 and SEC23B, to associate with β1-integrins. We found that GFP-RAB10, GFP-RAB13 and GFP-SEC23B readily associate with endogenous β1-integrin (Fig. 6A, S3I). In cells, GFP-RAB13 localized clearly on large β1-integrin endosomal vesicles and overlapped with β1-integrin on the cell surface (Fig. 6B). In contrast, enrichment of RAB10 and SEC23B to integrin-positive vesicles was less apparent and they did not overlap with β1-integrin on the plasma membrane (Fig. 6B; S3J). Furthermore, endogenous RAB13 and β1-integrin overlapped at steady state in MDA-MB-231 cells (Fig. 6C) and both RAB13 and β1-integrin associated with GFP-GGA2, alongside known GGA2 interactors clathrin and GGA3, which were also detected in the pulldown (Fig. 6D). Taken together, RAB10, RAB13 and SEC23B represent previously unknown GGA2 proximal proteins that associate with β1-integrins. In addition, RAB13, in particular, exhibits significant overlap with β1-integrins in intracellular vesicles as well as on the plasma membrane.

**Figure 6.**
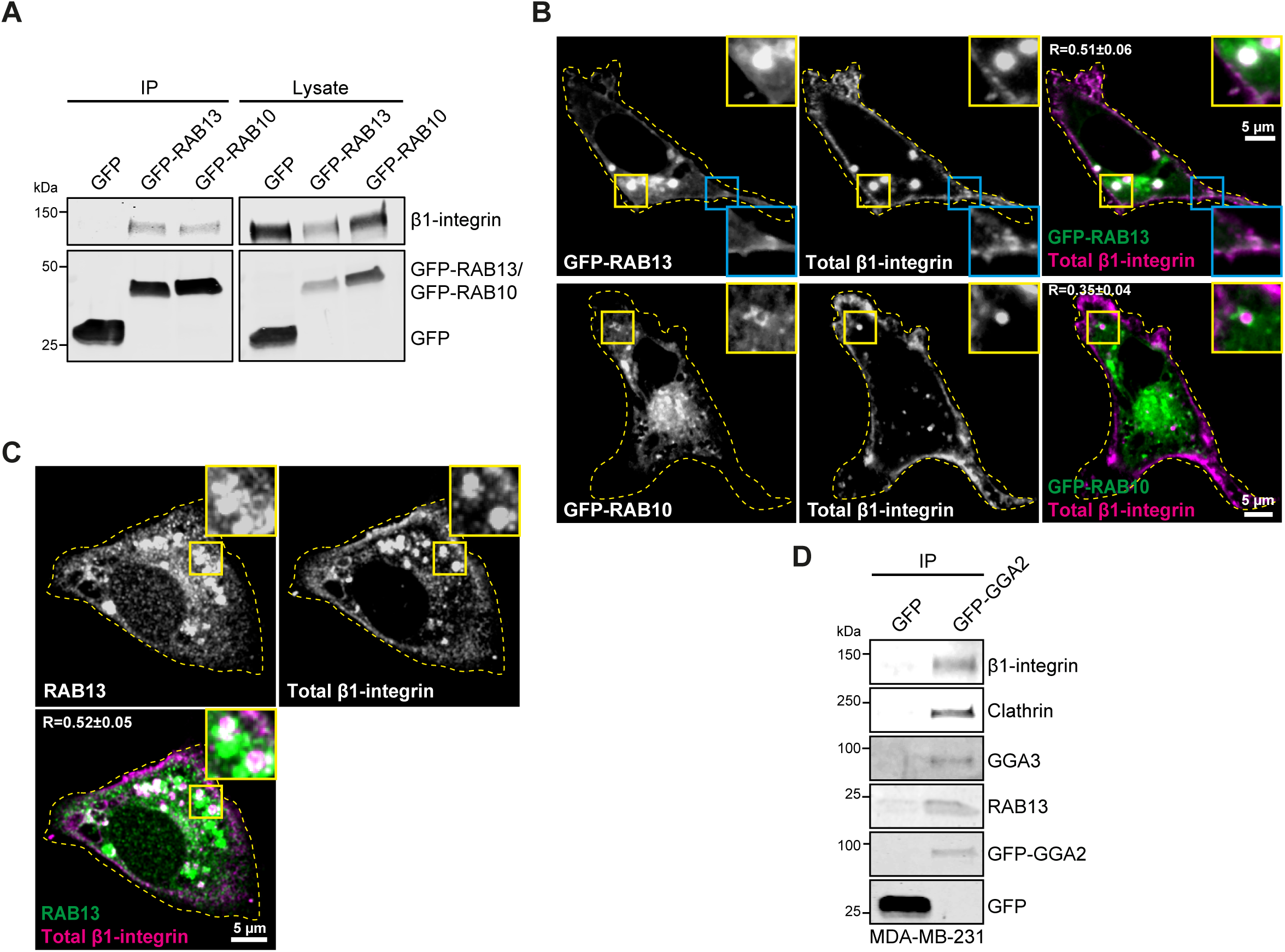
RAB13 and RAB10 associate with β1-integrin and GGA proteins. **A)** Representative western blots of GFP-Trap pulldowns in MDA-MB-231 cells transiently transfected with GFP, GFP-RAB10 or GFP-RAB13 and blotted for GFP and endogenous β1-integrin (n = 3 independent experiments). **B)** Representative confocal microscopy images of MDA-MB-231 cells transfected with GFP-RAB13 or GFP-RAB10 and stained with anti-β1-integrin antibody P5D2. Cytosolic overlap is highlighted in yellow insets and overlap on the membrane in blue insets. The extent of membrane overlap between the indicated proteins is quantified (R ± SD, n = 60-100 cells per condition). **C)** Representative confocal microscopy images of MDA-MB-231 cells stained for endogenous RAB13 and β1-integrin. Overlap between the indicated proteins is quantified (R ± SD, n = 150 cells per condition). Yellow insets show ROI. **D)** Representative western blots of GFP-Trap pulldowns in MDA-MB-231 cells transiently transfected with GFP or GFP-GGA2 and analyzed for GFP, β1-integrin and RAB13 and known GGA2-interacting proteins (GGA3 and clathrin) as positive controls.

### RAB13 is required for recycling of active β1-integrin

To assess if RAB10, RAB13 or SEC23B are involved in trafficking of β1-integrins, we first repeated the biotin-IP-based β1-integrin endocytosis assay (Fig. 1A) in cells silenced for RAB10, RAB13 or SEC23B. Depletion of RAB13 (siRNA#1) significantly increased the level of intracellular β1-integrin following 20 minutes of endocytosis (Fig. 7A,B; Fig. S4A), similarly to GGA2 silencing (Fig. 1A). In contrast, silencing of RAB10 or SEC23B had no significant effect on β1-integrin traffic (Fig. S4B-F), suggesting that these proteins may regulate other GGA2-dependent processes than β1-integrin traffic. Importantly, silencing of GGA2, RAB10 or RAB13 proteins did not influence the protein levels of the other GGA2-proximal components (Fig. S4A,D,G,H) indicating that their expression/stability are independently regulated.

**Figure 7.**
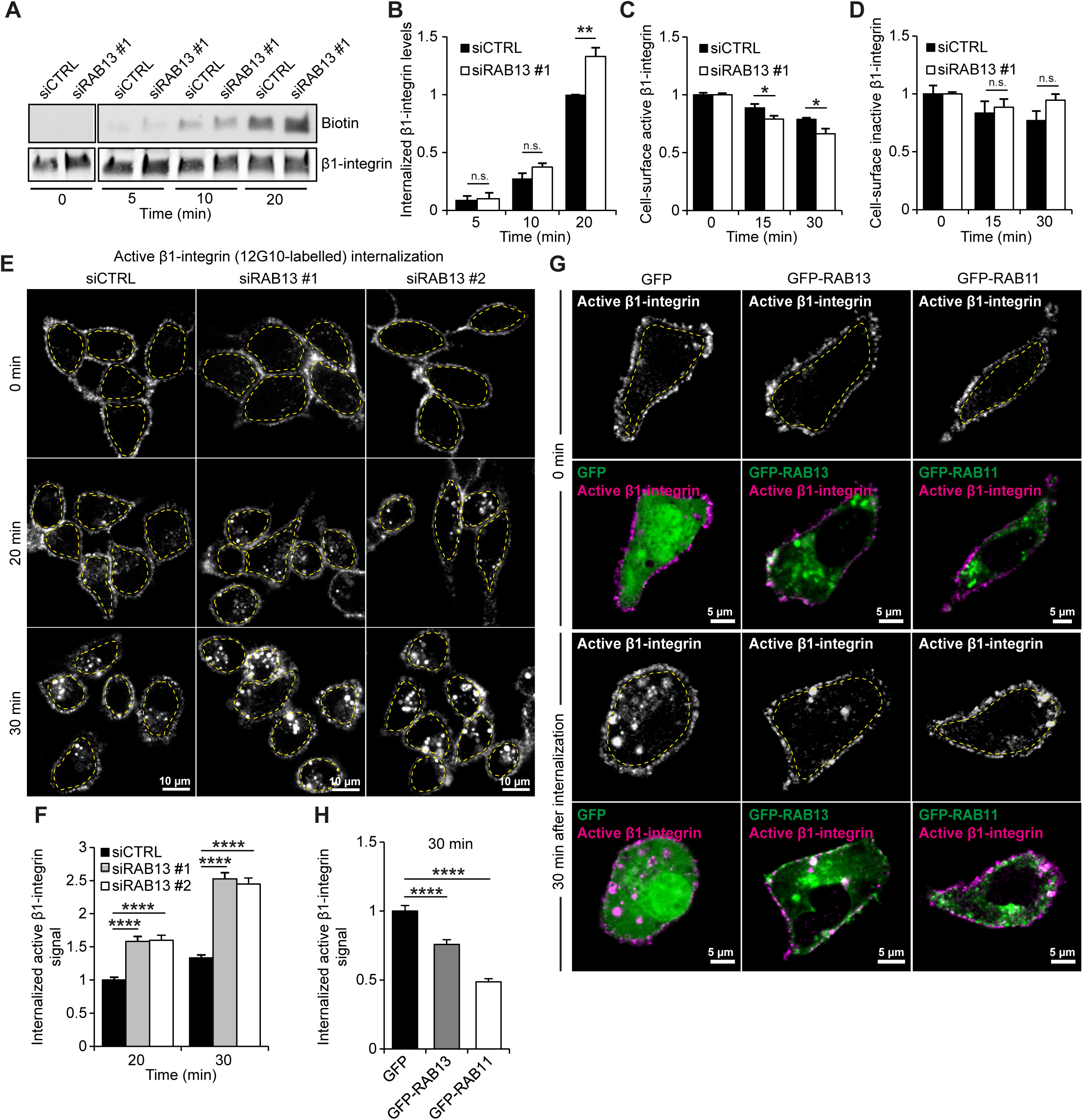
RAB13 supports recycling of active β1-integrin. **A-B)** Internalization of biotinylated cell-surface β1-integrin in siCTRL or siRAB13#1 MDA-MB-231 cells. Shown is a representative western blot and quantification of biotinylated β1-integrin relative to total β1-integrin and normalized against siCTRL 20 min time point (data are mean ± SEM of n = 3 independent experiments). **C-D)** Flow cytometry analysis of cell-surface levels of active (12G10 antibody) (C) or inactive (MAB13 antibody) β1-integrin (D) in siCTRL or siRAB13#1 silenced MDA-MB-231 cells following the indicated internalization periods, normalized to siCTRL at 0 min time point (data are mean ± SEM of n = 3 independent experiments). **E-F)** Representative confocal microscopy images (E) and quantification (F, performed as in 1L) of internalized active β1-integrin (12G10 antibody) at the indicated time points in control siRNA or RAB13 siRNA#1 or RAB13 siRNA#2 silenced MDA-MB-231 cells (data are mean ± SEM.; n = 100-120 cells per condition). **G-H)** Representative confocal microscopy images (G) and quantification (H, performed as in 1L) of internalized active β1-integrin (12G10 antibody) at the indicated time points in GFP GFP-RAB13 and GFP-RAB11 transiently transfected MDA-MB-231 cells (data are mean ± SEM.; n = 90-130 cells per condition). Statistical analysis for quantifications (n.s.= not significant, *p < 0.05, **p < 0.005, ***p < 0.0005. student’s unpaired *t*-test).

Congruently with the results obtained with GGA2 depletion (Fig. 1-2), silencing of RAB13 significantly increased the uptake of cell-surface-labelled active, but not inactive, β1-integrins in the flow cytometry based internalization assay (Fig. 7C,D). These data were further validated by silencing RAB13 with two independent siRNA and analyzing active and inactive β1-integrin internalization from the cell surface using immunofluorescence. Loss of RAB13 resulted in a significant increase in the accumulation of active β1-integrin in intracellular vesicles (Fig. 7E,F). Conversely, overexpression of GFP-RAB13 significantly reduced active β1-integrin accumulation inside cells (Fig. 7 G,H). In line with GGA2 overexpression (Fig. S1E, F), GFP-RAB13 overexpression had no significant effect on the localization of cell-surface-labelled inactive β1-integrin (Fig. S4I, J). Inhibition of receptor recycling with primaquine abolished the effect of GFP-RAB13 expression on active β1-integrin internalization (Fig. S4K,L), indicating that RAB13 influences recycling of active β1-integrin in these cells. Interestingly, the ability of GFP-RAB13 to support integrin recycling was similar to GFP-RAB11 (Fig. 7G, H), a well-established regulator of integrin recycling (Bridgewater et al., 2012). One of the other GGA2-proximal proteins identified in the BioID was Arf6, which has been implicated both in integrin endocytosis and in integrin recycling (Powelka et al., 2004; Fang et al., 2010; Schweitzer et al., 2011; Dunphy et al., 2006; Sakurai et al., 2010; Chen et al., 2014). Interestingly, we found that while loss of RAB13 significantly increases intracellular accumulation of β1-integrin, ARF6 silencing significantly reduces integrin intracellular uptake (Fig. 8A-C). These data indicate that either ARF6 is not involved in the GGA2-RAB13-mediated integrin recycling pathway, or that the reciprocal roles of ARF6 in integrin recycling and endocytosis complicates the interpretation of results in ARF6-silenced cells, and to fully distinguish ARF6 function in integrin recycling would require an acute method of ARF6 depletion following integrin internalization.

**Figure 8.**
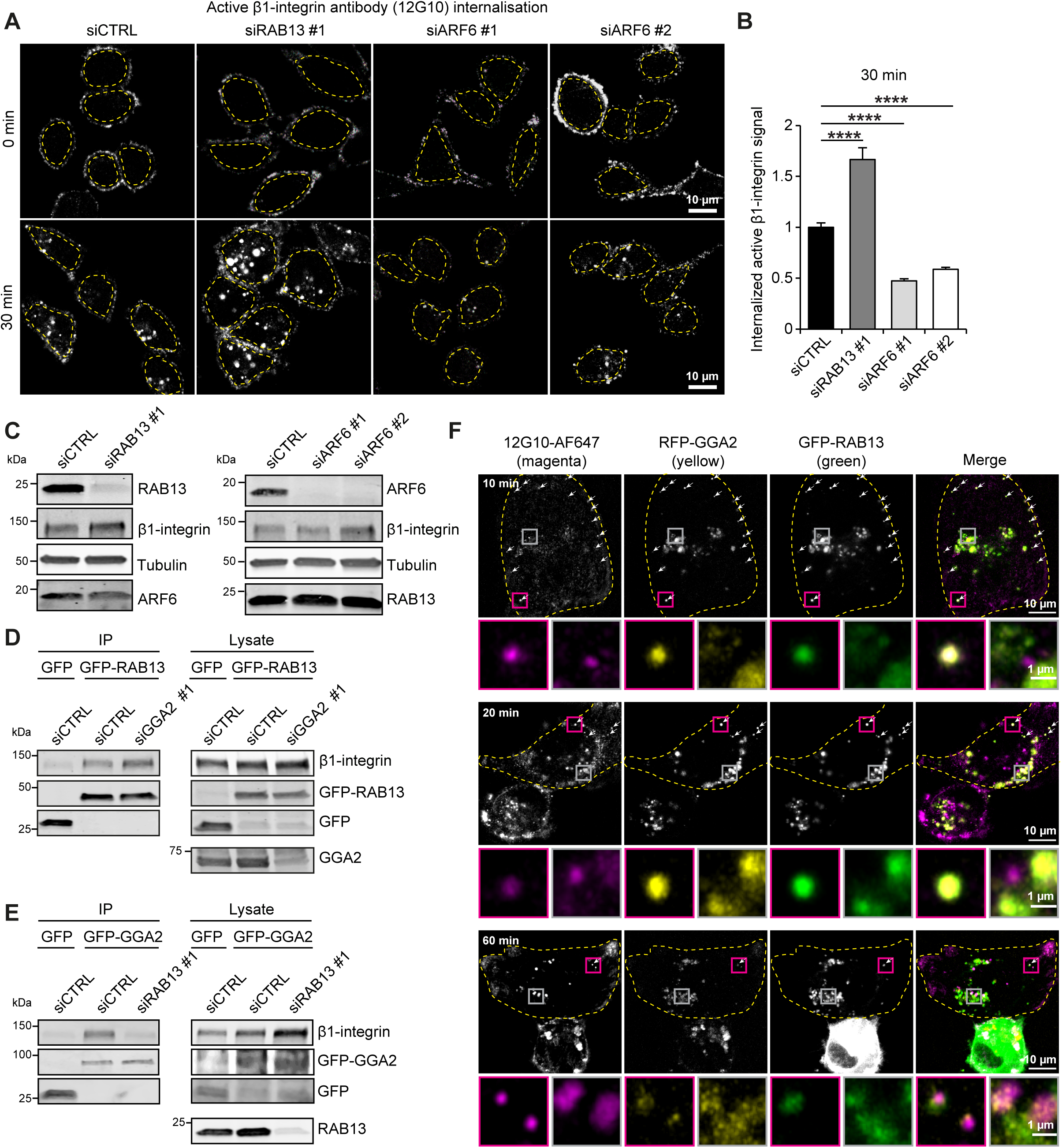
GGA2 association with β1-integrin is RAB13 dependent. **A-B)** Representative confocal microscopy images (A) and quantification (B, performed as in 1L) of internalized active β1-integrin (12G10 antibody) at the indicated time points in control siRNA or siRAB13#1 or ARF6 siRNA#1 or ARF6 siRNA#2 silenced MDA-MB-231 cells (data are mean ± SEM from n = 100-120 cells) **C)** Representative western blot analysis of protein expression in MDA-MB-231 cells transiently transfected with siRAB13#1 or siARF6#1 or siARF6#2. **D)** Representative western blot analysis of GFP-Trap pulldowns in MDA-MB-231 cells transiently transfected with GFP or GFP-RAB13 and silenced with control siRNA or GGA2 siRNA. **E)** Representative western blot analysis of GFP-Trap pulldowns in MDA-MB-231 cells transiently transfected with GFP or GFP-GGA2 and silenced with control siRNA or RAB13 siRNA. **F)** Dynamic colocalization of GGA2, RAB13 and β1-integrin. Cells were transfected with RFP-GGA2, GFP-RAB13 and plated on fibronectin-coated glass-bottom dishes. Alexa Fluor 647-labeled antibody recognizing the active integrin conformation (12G10-AF647) was added to the media at 3.7 µg/ml and cells were fixed after 10, 20 or 60 min. The images were acquired in Airyscan super-resolution mode and show are maximum projections of the cell. The magenta insets show the situation at the cell periphery and the grey insets show the perinuclear region. Arrows highlight endosomes where all three proteins overlap. The dashed line represents the cell outline.

Our data thus far indicate that GGA2 and RAB13 have similar effects on integrin traffic and can associate with β1-integrin and with each other. Therefore, we were interested to investigate how RAB13 and GGA2 are recruited to β1-integrin. We performed GFP-trap pulldowns of GFP-GGA2 or GFP-RAB13 in control silenced cells or cells silenced for the respective partner (RAB13 for GGA2 pulldowns and vice versa). Silencing of GGA2 did not affect β1-integrin association with GFP-RAB13 (Fig. 8D). In contrast, RAB13 silencing attenuated β1-integrin association with GFP-GGA2 (Fig. 8E), indicating that GGA2 associates with β1-integrin in a RAB13-dependent manner.

To assess the overlap of GGA2, RAB13 and β1-integrin in cells over time, we imaged the trafficking dynamics of an Alexa Fluor 647-labeled antibody recognizing the active integrin conformation (12G10-AF647) in cells transfected with RFP-GGA2 and GFP-RAB13. The three proteins overlapped 10 minutes post antibody addition, in small vesicles at the cell periphery (Fig. 8F). GGA2 and RAB13 overlapped predominantly with each other in the perinuclear region, but once β1-integrin accumulated in bright endosomes in the perinuclear region (60 min), it no longer colocalized with GGA2 and RAB13 (Fig. 8F, bottom panel). Taken together, RAB13 and GGA2 overlap dynamically with trafficking β1-integrin in the cells and silencing of RAB13 and GGA2 significantly impairs cell-surface targeting of endocytosed active β1-integrin indicating a joint or sequential regulation of active receptor dynamics by these proteins.

### GGA2 and RAB13 are required for efficient cancer cell migration and invasion

The β1-integrin recycling defect in RAB13-depleted cells resembled the situation in GGA2 knockdown. Furthermore, RAB13 has been shown to support cancer cell invasion (Ioannou et al., 2015). Thus, we decided to investigate whether RAB13 reduces the amount of active β1-integrin in focal adhesions similarly to GGA2. RAB13 silencing had no significant effect on focal adhesion number 30 or 60 minutes after plating but significantly counteracted the decline of focal adhesion numbers detected in the control cells after 2 hours (Fig. 9A). The relevance of this needs further investigation. In addition, RAB13 significantly reduced the levels of active β1-integrin in the adhesions at all time points after plating (Fig. 9B,C). Next, we investigated whether the reduced migration of GGA2-knockdown cells is also reproduced in RAB13 knockdown. On fibronectin-coated plates, RAB13-depleted cells showed slower migration, similarly to GGA2-depleted cells, with all independent siRNA sequences tested (Fig. 9D-F, Video 1). RAB13 depletion appeared to also impact, albeit modestly, the directionality of cell migration on fibronectin (Fig. 9G, Video 1). The slower migration of cells was reflected also in the somewhat reduced membrane protrusion dynamics of GGA2-and RAB13-depleted cells (Fig. 9H, I, Video 2). Importantly, these data were also relevant for cancer cell invasion in vivo in a zebrafish embryo xenograft model, where depletion of GGA2 or RAB13 significantly reduced cancer cell invasion (Fig. 9J). These in vivo effects were specifically due to reduced invasion since GGA2 or RAB13 silencing did not influence cell viability of zebrafish embryo xenografts (Fig. S4M). Taken together, these results show that interfering with the GGA2 and RAB13-dependent delivery of β1-integrins to the plasma membrane leads to reduced cell motility and impaired cancer cell invasion in vitro and in vivo.

**Figure 9.**
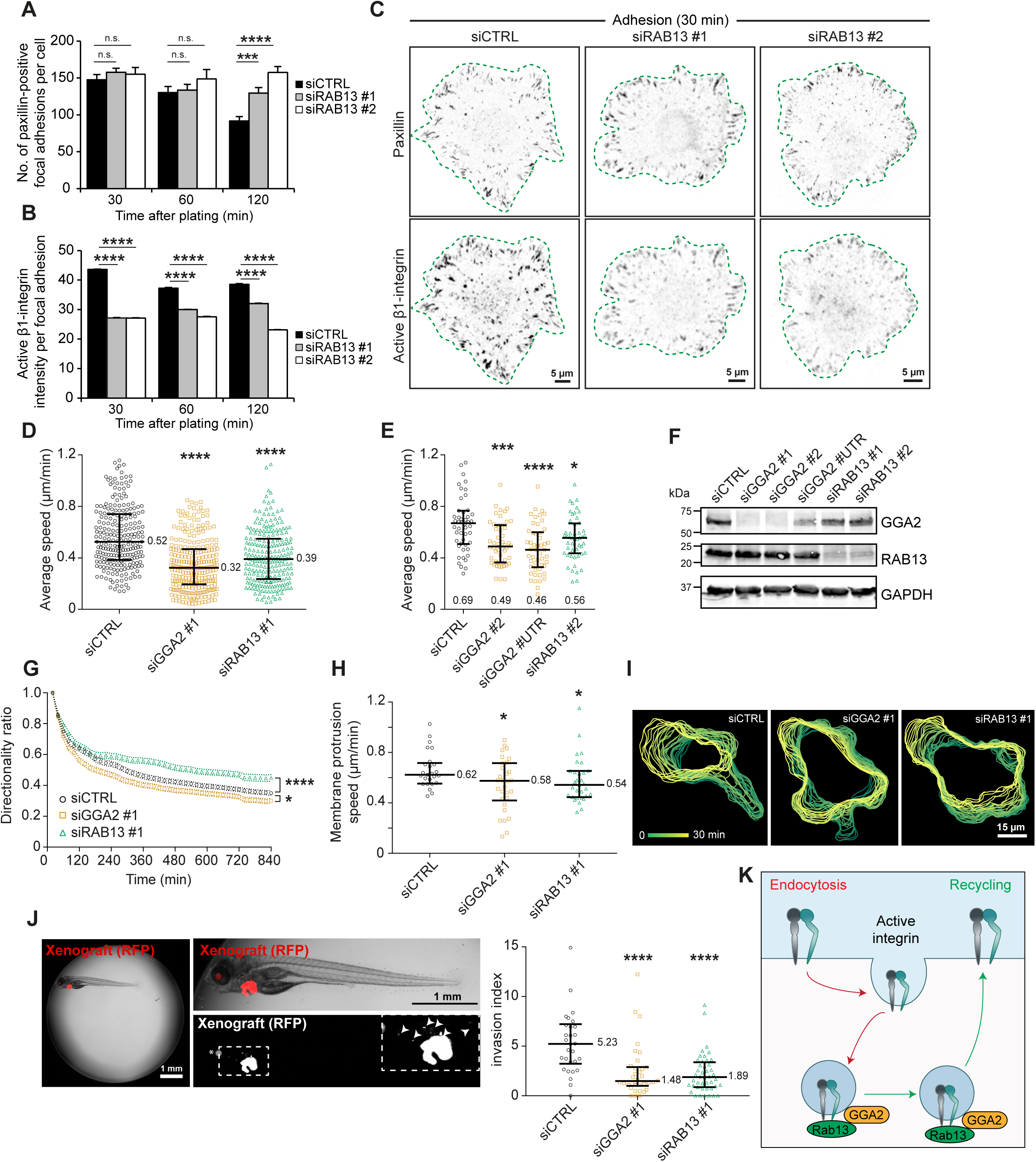
GGA2 and RAB13 support efficient cancer cell migration and invasion. **A-C)** siCTRL, siRAB13 #1 or siRAB13 #2 MDA-MB-231 cells were plated on collagen-fibronectin for the indicated time points, fixed and stained for paxillin (to detect focal adhesions) and active β1-integrin (12G10). Quantifications of focal adhesion number (A) and 12G10 mean fluorescence intensity in focal adhesions (B) and representative images of cells (bottom plane) (C) are shown. Data are mean ± SEM; n = 7,500-16,300 focal adhesions from 80-105 cells per condition. **D-E)** Average speed of single cells during 14 h of 2D migration on fibronectin-coated dishes. GGA2 or RAB13 were depleted by different siRNAs for 3 days. In all panels, bars represent median and interquartile range and median values are indicated. P-values in all following figure panels result from testing the difference between control and treated samples by unpaired Mann-Whitney test (n = 230 to 264 cells from three independent experiments per condition (D); n = 49 to 50 cells from one experiment (E). Two outlier data points in siCTRL condition are not displayed in (E) for clarity. **F)** GGA2 and RAB13 expression in cell lysates from siRNA-treated cells used for the migration experiment in (D-E). GAPDH was used as a loading control. **G)** Directionality ratio of cell trajectories from (D) over time. Directionality ratio was calculated as (start-finish distance/total trajectory length). The arithmetic mean of all tracks per condition is shown. Error bars represent SEM. Final directionality ratios: siCTRL = 0.36, siGGA2 #1 = 0.32, siRAB13 #1 = 0.46. **H)** Membrane protrusion speed of cells migrating on fibronectin-coated dishes. MDA-MB-231 cells expressing Lifeact-mRFP were imaged once every 1 min. Segmented cell outlines from 25 to 31 cells from two independent experiments over a 30 min time interval were used for quantification. **I)** Example of segmented cell outlines over a 30 min time interval. **J)** Invasion of siCTRL, siGGA2 #1 or siRAB13 #1 MDA-MB-231 cells in zebrafish embryos. Cells expressing GFP or RFP were transplanted into the pericardial cavity of 2-day old zebrafish embryos and after 4 days the invasion index of the tumors was measured. Example pictures are from siCTRL MDA-MB-231 tumor xenografts expressing RFP. Magnified region is outlined with a dashed line, invading tumor cells are labelled with arrow heads and non-specific fluorescence in the lens with an asterisk (*) Data points represent individual embryos. Invasion index in 30 to 44 embryos from two independent experiments was quantified as number of invading cells normalized to tumor fluorescence intensity. One outlier data point in siCTRL condition is not displayed. **K)** Graphical summary of GGA2 and RAB13 involvement in β1-integrin traffic.

Currently, it is unclear whether the GGA2/RAB13-mediated active β1 integrin traffic plays a functional role in cancer metastasis. However, we find that in comparison to normal tissue, GGA2 (Fig. 10A) and RAB13 (Fig. 10B) mRNA expression levels are significantly upregulated in some breast tumor subtypes. Thus, RAB13-and GGA2-driven cell motility and invasion may also play a role in human breast cancer progression. However, this needs to be thoroughly investigated.

**Figure 10.**
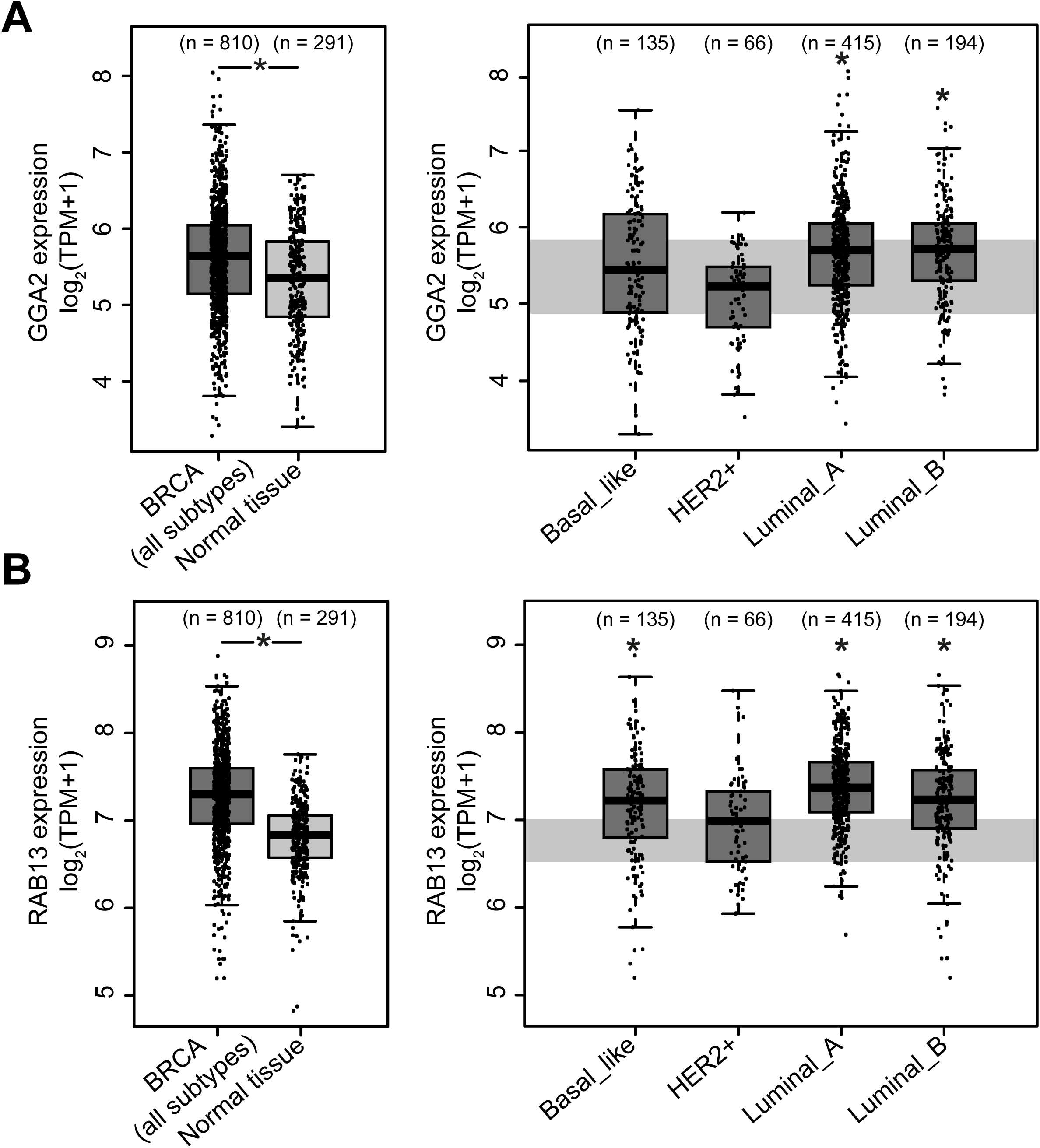
GGA2 and RAB13 are highly expressed in some subtypes of breast cancer. **A-B)** Differential analysis of GGA2 (A) and RAB13 (B) mRNA expression in TCGA tumor samples (dark grey boxplots) and in paired adjacent TCGA and GTEx normal samples (light grey boxplots) for all breast cancer subtypes (left-hand panels) and for specific breast cancer subtypes (right-hand panels). Data were compiled using the GEPIA2 platform (see methods). The data from normal samples in left-hand panels are shaded in light grey in the right-hand panels. The boxplot shows median and interquartile range; method for differential gene expression analysis, one-way ANOVA; p-value < 0.05; TPM: transcripts per million.

## Discussion

β1-integrin trafficking plays an important role in the dynamic regulation of cell adhesion and motility and is implicated in malignant processes such as cancer cell invasion (Bridgewater et al., 2012; De Franceschi et al., 2015). However, the molecular mechanisms and proteins regulating the specificity of integrin traffic remain poorly characterized. Here, we have used RNAi screening and BioID to identify new regulators of integrin traffic and demonstrate a role for GGA2, and its previously unknown proximal binding partner RAB13, in the trafficking of active β1-integrins (Fig. 9K).

Our work suggests GGA2 as an important determinant of activity-dependent integrin traffic and hence cancer-related processes, such as cell migration and invasion. We reported previously that GGA3-depletion results in lysosomal targeting and downregulation of collagen binding α2β1-integrin, due to mislocalization of the β1-integrin recycling component SNX17 (Steinberg et al., 2012; Böttcher et al., 2012) to late endosomes (Ratcliffe et al., 2016). In contrast, GGA2 is unique in its integrin regulation as it does not affect β1-integrin stability or levels, but instead specifically facilitates the trafficking of active β1-integrins. Increased active β1-integrin accumulation in GGA2-silenced cells is fully reversed upon treatment with the membrane recycling inhibitor primaquine, implicating a role of GGA2 in the recycling of active β1-integrin to the plasma membrane.

A role for GGA2 in mediating receptor recycling is surprising since GGA2 has been described to localize to the trans-Golgi network and clathrin-coated vesicles (Dell’Angelica et al., 2000; Puertollano et al., 2001). In addition, a “knocksideways”-depletion indicates a role for GGA2 in anterograde but not retrograde Golgi trafficking (Hirst et al., 2012). However, the biological role and subcellular localization of GGA2 has not been studied extensively in carcinoma cells. Focusing on invasive breast cancer MDA-MB-231 cells, where integrin traffic is implicated in the regulation of cell motility (Gu et al., 2011; Mai et al., 2011; Pellinen et al., 2006), we found a subpopulation of GGA2 partially overlapping with early and recycling endosomal markers.

Furthermore, we employed BioID proximity biotinylation (Roux et al., 2012) to define GGA2 proximal proteins in situ in MDA-MB-231 cells. We identified two RAB family small GTPases, RAB13 and RAB10 as well as the vesicle coat protein SEC23B, associating with both GGA2 and β1-integrin. Loss-of-function experiments validated a role for RAB13 in the recycling of active β1-integrin. RAB10 and SEC23B, on the other hand, may regulate some other GGA2 and integrin functions, but this remains to be investigated. The lack of an integrin trafficking phenotype following SEC23B knockdown could also be explained by the unperturbed expression and function of the paralog SEC23A (Khoriaty, PNAS 2018). We identified other known (e.g., IGFR2), as well as new, GGA2-proximal proteins (Table S1). In our list of interactors, we only display candidates scoring positive in three or more experiments. However, notably, three of the proteins affected by knocksideways depletion of GGA2 in HeLa cells (RABEP1, TNFAIP8 and PARP4) (Hirst et al., 2012) were also identified in our BioID, but only in 2 out of 4 assays. Thus, while not all of the identified GGA2 interactors are likely to regulate integrin traffic, they may be vital GGA2 effectors in other yet unknown biological processes.

RABs play important roles in different steps of receptor trafficking and are linked to cancer progression (Subramani, Alahari 2010). Two specific RABs, RAB21 and RAB25, have previously been shown to regulate integrin traffic by directly binding to integrins. RAB21 is widely expressed in different cell types. It recognizes a conserved motif found in the α-tails of all integrins (apart from α11) and regulates integrin endocytosis (Pellinen et al., 2006; Mai et al., 2011). RAB25, on the other hand, is expressed only in specific epithelial tissues and some carcinomas (Wang et al., 2017) and associates with the fibronectin-binding α5β1-integrin to drive its recycling (Caswell et al., 2007). Previous studies have shown the role of RAB13 in trafficking of receptors between trans-Golgi network and transferrin-positive recycling endosomes in epithelial cells (Nokes et al., 2008) and activation of RAB13 by guanine nucleotide exchange factor DENND2B on the plasma membrane has been linked to cancer cell migration and invasion (Ioannou et al., 2015). We find that RAB13 silencing impairs active integrin traffic similarly to GGA2 depletion. Moreover, RAB13 also colocalizes with GGA2 and β1-integrin in endosomes and the plasma membrane, suggesting that the previously described RAB13-mediated defects in cancer cell migration and invasion (Ioannou et al., 2015) may involve the GGA2 and β1-integrin complex.

RAB GTPases are the master regulators of intracellular membrane traffic and are involved in virtually all steps of trafficking including vesicle formation, transport and fusion. There are still many unknown functions for RABs in membrane trafficking and in human ailments. Furthermore, the role of trafficking adaptors, such as GGAs, is not fully explored in human cancer. Recently, several multiprotein complexes have been assigned as regulators of integrin traffic, including the retriever complex (McNally et al., 2017) and the CORVET complex (Jonker et al., 2018). Our data, putting forward a role for a β1-integrin, GGA2 and RAB13 complex in the recycling of active β1-integrins contributes to the view of multi-subunit complexes regulating receptor traffic and contributing to the remarkable sophistication and specificity of membrane trafficking pathways that determine cellular fitness in a fundamental way.

## Acknowledgments

We thank P. Laasola and J. Siivonen for technical assistance, M. Saari for help with the microscopes. We also thank The Cell Imaging Core and Proteomics core facilities at Turku Centre for Biotechnology supported by the University of Turku and Biocenter Finland, for technical assistance. The Ivaska lab for critical reading and feedback on the manuscript. This study has been supported by the Academy of Finland (J.Ivaska), Academy of Finland CoE for Translational Cancer Research (J.Ivaska), ERC CoG grant 615258, Sigrid Juselius Foundation, Orion Research Foundation and the Finnish Cancer Organization (J.Ivaska). P.S. has been supported by the Turku Doctoral Program of Molecular medicine (TuDMM). J.Icha is a member of the Turku Collegium of Science and Medicine and recipient of the EMBO Long-Term Fellowship ALTF 405-2018.

## Authors contributions

Conceptualization, J.Ivaska.; Methodology, P.S., J.A., A.A., I.P., J.Icha, J.Ivaska.; Formal Analysis,.P.S., A.R., A.A., I.P., J. Icha; Investigation, P.S., J.A., A.A., I.P., M.P., J. Icha, J.Ivaska.; Writing– Original Draft, J.Ivaska, P.S. and H.H.; Writing – Review and Editing, J.I., P.S., A.R., J.A, H.H., J. Icha; Visualization, P.S., J.A., I.P., J. Icha, H.H. Supervision, J.I.; Funding Acquisition, J.I.

## Video legends

### Video 1

Migration of cells in 2D is affected by GGA2 or RAB13 knockdown. Cells transfected with siRNA for 3 days were plated on fibronectin-coated 24-well plates. Single plane phase contrast 2×2 tiled image was acquired every 20 min for 14 hours. Manual tracking trajectories are overlaid. Scale bar: 200 µm.

### Video 2

Lamellipodial dynamics of MDA-MB-231 cells migrating on fibronectin. Filamentous actin was labelled by stable expression of Lifeact-mRFP. The bottom plane of the cells was imaged once per minute. The control and siRNA-treated cells were imaged simultaneously in the same plate. Scale bar: 15 µm.

## Materials and methods

### RNAi Screening

Cell Spot Microarrays (CSMA) were conducted by printing matrigel, siRNA and lipid transfection reagent containing spots as arrays, followed by cell plating to silence target genes (Rantala et al., 2011a). GGA family member and interactor targeting siRNAs (Oligo IDs indicated in Fig. S1A) and control siRNA were purchased from Qiagen. Before plating MDA-MB-231 cells on CSMA, the arrays were blocked with Lutrol F108 (Univar AB) for 15 min at room temperature. Cells were washed and harvested using HyQTase (HyClone, SV30030.01), then plated on CSMA and endocytosis was measured 72 h after the reverse siRNA transfection using an antibody-based β1-integrin endocytosis assay described previously (Arjonen et al., 2012). In short, non-function blocking β1-integrin antibody (K20) was labelled using an Alexa Fluor 488 protein labelling kit (Invitrogen). Cells plated in the CSMA were then incubated live with the K20-Alexa Fluor 488 conjugated antibody for 1 h on ice to allow labelling of cell-surface integrins. The CSMA was then washed and endocytosis of cell-surface labelled β1-integrins triggered by incubating cells at 37°C for 30 min. Any residual cell-surface fluorescence was then quenched using anti-Alexa Fluor 488 antibody (Invitrogen) for 1 h on ice and cells were then fixed using 4% paraformaldehyde, permeabilized and incubated with anti-mouse Alexa Fluor 555 secondary antibody to label both internal and cell surface K20 pools (total β1-integrin). Levels of detected endocytosed β1-integrin were normalized against total β1-integrin levels. Nuclei were labelled using Syto60 (Invitrogen). Fluorescence intensities were measured using Laser microarray scanner (Tecan LS400) and ArrayPro Analyzer v4.5 software.

### Cell culture

MDA-MB-231 cells (ATCC) were grown in Dulbecco’s modified essential medium (DMEM; Sigma-Aldrich, D5769) supplemented with 10% fetal bovine serum (FBS; Sigma-Aldrich, F7524), 1% penicillin/streptomycin (Sigma-Aldrich, P0781-100ML), 1% MEM non-essential amino acids (Sigma-Aldrich, M7145) and 1% L-glutamine (Sigma-Aldrich, G7513-100ml). Cells were regularly tested for mycoplasma infection and were grown at +37°C, 5% CO_2_, until 70–80 % confluence before washing with PBS (Sigma-Aldrich, D8537), detaching with trypsin (Sigma-Aldrich, T4174-100ML) and plating onto new dishes.

### Transient transfections

DNA construct transfections were carried out using Lipofectamine 3000 (Invitrogen, L3000-015) or jetPRIME reagent (Polyplus transfection). siRNA was transfected using Lipofectamine RNAiMAX (Invitrogen, 13778-150) according to the manufacturer’s instructions. Cells were transfected with DNA constructs for 24 h or with siRNAs for 72 h before the experiment. The siRNAs used in this study are indicated in the reagent table. For rescue experiments, cells were transfected with siRNA for 72 h and at the end of 48 h, transfected with the DNA construct.

### Recombinant DNA

RFP-GGA2, GFP-GGA2 and myc-BioID-GGA2 constructs were generated by cloning Human GGA2 from GGA2 ORF entry vector (RC200153, origene) into following vectors using cloneEZ: pTagRFP-c (FP141, Evrogen), pTagGFP2-C (FP191, Evrogen) and pcDNA3.1 mycBioID (gift from Kyle Roux (Roux et al., 2012) (Addgene plasmid # 35700; RRID:Addgene_35700). GFP-RAB10, GFP-RAB13 and GFP-SEC23B constructs were generated by the Genome Biology Unit cloning service (Biocenter Finland, University of Helsinki). Briefly, GFP entry clone from the human ORFeome collaboration library (RAB10, 100066678; RAB13, 100002881; SEC23B, 100003783) was transferred into the pcDNA6.2/N-emGFP destination vector using the standard LR reaction protocol. Other constructs included RFP-RAB5 (described in (De Franceschi et al., 2016)), DsRed-RAB7 (Addgene plasmid #12661), GFP-RAB4 and GFP-RAB11 (described in (Arjonen et al., 2012)) and GFP-EEA1 (Addgene plasmid #42307).

### Western blot analysis

Reduced and heat-denatured protein extracts were separated by SDS-PAGE (4–20% Mini-4– 20% Mini-PROTEAN® TGX™ Gel, Bio-Rad, 456-1094) and then transferred onto 0.2 µm nitrocellulose membranes (Trans-Blot Turbo Transfer Pack; Bio-Rad, 170-4158). Membranes were blocked with blocking buffer (Thermo Scientific™ Pierce™ StartingBlock™; 10108313) for 30 min at room temperature and incubated with primary antibodies (see key resources table) diluted in blocking buffer overnight at +4°C with agitation. On the following day, membranes were washed three times with TBST for 10 min each and incubated with the appropriate fluorophore-conjugated secondary antibodies (see key resources table) (1:5000, diluted in blocking buffer) for 1 h at room temperature. Finally, membranes were scanned using the Odyssey (LI-COR) imaging system.

### Biotin-based internalization assay

Cells in 6 cm dishes were placed on ice and washed with cold PBS. Cell-surface receptors were labelled with 0.5 mg/ml cleavable biotin on ice for 30 min (EZlink cleavable sulfo-NHS-SS-biotin; Thermo Scientific, 21331) in Hanks’ balanced salt solution (Sigma, H9269). Unbound biotin was washed away with cold PBS and internalization of receptors was triggered by addition of pre-warmed serum-free media and incubation at 37°C for the indicated time points with the exception of the 0 min time point. In this case, cold media was added to the 0 min dishes and cells were kept on ice to prevent internalization of receptors. For primaquine treatment, primaquine was diluted in PBS and added both to the biotin labelling mix and to the pre-warmed media at a final concentration of 100 µM. Following internalization, for the indicated time points, cells were transferred to ice and surface biotin was cleaved with 60 mM MesNa (Sigma, 63705) in MesNa buffer (50 mM Tris-HCl [pH 8.6], 100 mM NaCl) on ice for 30 min, followed by quenching with 100 mM iodoacetamide (IAA, Sigma, I1149-25G) on ice for 15 min. Cells were washed with cold PBS, scraped in 300 µl lysis buffer (50 mM Tris pH 7.5, 1.5 % Triton X-100, 100 mM NaCl, protease (Sigma, 4906837001) and phosphatase (Sigma, 5056489001) inhibitor tablets) and incubated at 4°C, with agitation, for 20 min. Afterward, the lysates were clarified by centrifugation (10,000×g, 10 min, 4°C), and incubated overnight with a β1-integrin antibody, P5D2, for immunoprecipitation. Lysates were then incubated with protein G sepharose beads (GE Healthcare, 17-0618-01) for 2–3 h at 4°C and beads were washed 3 times with diluted lysis buffer (dilution of 1:3 in dH2O) followed by elution in non-reducing Laemmli sample buffer and denaturation for 5 min, 95°C. Western blotting was performed as described above and biotinylated (internalized) β1-integrin was detected by immunoblotting with horseradish peroxidase (HRP)-conjugated anti-biotin antibody (Cell Signaling Technology, 7075, 1:15000) and imaging with the chemiluminescence film-based system. Total β1-integrin was detected by immunoblotting the membrane again with clone N29 anti-β1-integrin antibody (Merck, MAB2252, 1:1000) and appropriate secondary antibody and analysis on the Odyssey (LI-COR) imaging system. Biotin and total β1-integrin signals were quantified as integrated densities of protein bands with ImageJ (v. 1.43u), and the biotin signal was normalized to the corresponding total β1-integrin signal for each time point.

### Co-Immunoprecipitation

Cells expressing GFP-conjugated proteins (one 10 cm dish per condition) were washed with cold PBS and detached with trypsin. Following centrifugation, 200 μl of IP-lysis buffer (40 mM HEPES-NaOH, 75 mM NaCl, 2 mM EDTA, 1% NP-40, protease and phosphatase inhibitor tablets) was added to the cell pellet and cells were lysed at +4 °C for 30 min, followed by centrifugation (10,000×g for 10 min, +4 °C). 20 μl of the supernatant was kept aside as the lysis control. The remainder of the supernatant was incubated with GFP-Trap beads (ChromoTek; gtak-20) for 1 h at 4°C. Finally, immunoprecipitated complexes were washed three times with wash buffer (20 mM Tris-HCl (pH 7.5), 150 mM NaCl, 1 % NP-40) followed by denaturing for 5 min at 95°C in reducing Laemmli buffer before analyses by SDS-PAGE followed by immunoblotting as above.

### Flow cytometry-based endocytosis assay

Attached cells were incubated with 1:500 diluted 12G10 or MAB13 antibodies in serum-free medium on ice for 30 min to allow the labelling of active and inactive cell-surface pools of β1-integrin. Unbound antibody was washed away in cold PBS and endocytosis was triggered by addition of pre-warmed serum-free medium to the cells and incubation at 37°C for 15 min and 30 min. In 0 min control cells, cold media was added and cells were kept on ice. Warm media was then removed and cells washed with cold PBS, detached with scraping the cells gently and fixed with 4% PFA in PBS for 15 min at RT. Cells were then washed with PBS and incubated with the appropriate Alexa Fluor-conjugated secondary antibody (dilution 1:300 in PBS) for 1 h at 4°C. Cells were finally washed with PBS and analyzed with LSRFortessa (BD Biosciences). Data analysis was done using Flowing software version 2 (Cell Imaging Core of the Turku Centre for Biotechnology; www. http://flowingsoftware.btk.fi/).

### Immunofluorescence, microscopy and image analysis

Cells were plated either on 8-well slides (µ-Slide 8 well, ibiTreat; Ibidi, 80826) or on 3.5 mm dishes (Ibidi ibiTreat 35mm; Ibidi, 80136). Cells were fixed in 4% paraformaldehyde for 15 min at room temperature, and following quenching of the fixative in 50 mM NH_4_Cl for 15 min at room temperature, cells were permeabilized with 30% horse serum in PBS containing 0.3% Triton X-100 for 10 min at room temperature. Afterward, cells were incubated with primary antibodies (see key resources table) diluted in 30% horse serum at +4°C, overnight. Later, cells were washed three times with PBS and appropriate secondary antibodies (see key resources table) diluted 1:250 in 30% horse serum were added along with DAPI (1:5000) for 1 h at room temperature followed by three PBS washes. Samples were stored in PBS protected from light at 4°C until imaged and were imaged at room temperature. Middle plane images were acquired using LSM780 or LSM880 (ZEISS) laser scanning confocal microscope or a Marianas spinning disk confocal microscope (3i) with a CSU-W1 scanning unit (Yokogawa). All the image analysis was done using FIJI (ImageJ version 1.49 k). Pearson’s correlation coefficient (R) of colocalization was calculated using the coloc2 plugin in FIJI. Number of focal adhesions were calculated on paxillin channel by background subtraction (rolling ball radius=20) and applying median Gaussian filter (3.0) followed by analyzing the number of particles represented as focal adhesions and saving them as an image mask. For measuring the β1-integrin intensity in focal adhesions, focal adhesion mask from the paxillin channel was superimposed on 12G10 channel and mean intensity of 12G10 in each focal adhesion was measured.

### ECM proteins and coating

Where indicated, wells were coated with 10 μg/mL collagen (PureCol® EZ Gel Bovine Collagen Solution, Type I; Advanced Biomatrix, 5074-G) or 10 μg/mL fibronectin (Merck-Millipore Calbiochem, 341631-5MG) in PBS for 1 h at 37°C. Unbound ligand (coating) was then removed, wells were blocked for 1 h at 37°C with 0.1% BSA and then washed with PBS before plating the cells in serum-free media.

### Microscopy-based endocytosis assay

Cells were plated on 35 mm ibidi dishes, washed with PBS and incubated with 12G10 or MAB13 antibodies in Hanks’ balanced salt solution on ice for 30 min to allow labelling of cell-surface active and inactive β1-integrin pools, respectively. Unbound antibody was washed away in cold PBS and endocytosis was triggered by addition of pre-warmed serum-free medium to the cells and incubation at 37°C for the indicated time points. In 0 min control cells, cold media was added and cells were kept on ice. The warm medium was then removed and cells washed with cold PBS, fixed, permeabilized, stained as described above and the middle plane of cells imaged with the LSM880 laser scanning confocal microscope (ZEISS) using the 63×/1.4 Plan-Apochromat oil objective. For quantification of internalized integrin signal, the area of interest was drawn inside the cell excluding the plasma membrane and mean integrated density was measured using ImageJ.

### Micropatterns and immunostaining

Micropatterns were created on glass coverslips (Azioune et al., 2009; Alanko et al., 2015). Glass coverslips were cleaned with 97% ethanol, dried with airflow, and exposed to deep UV light for 5 min. The clean coverslips were coated with 0.1 mg/ml poly(l-lysine)-graft-poly(ethylene glycol) (Surface Solutions) in 10 mM Hepes, pH 7.3, for 1 h at RT, rinsed once with PBS and once with water and dried. Crossbow-shaped 45-µm micropatterns were generated onto the pegylated coverslips using a photomask and exposing the coverslips to deep UV light for 6 min. The coverslips were washed with water, dried, and coated with 50 µg/ml fibronectin and 5 µg/ml fibrinogen Alexa Fluor 647 in PBS for 1 h at room temperature. Cells in serum-free medium were plated, left to adhere for 10 min at 37°C, washed with medium to remove non-adherent cells, and incubated for 3 h before fixing and staining as described above.

### Imaging of GGA2, RAB13 and β1-integrin colocalization

Cells transfected with RFP-GGA2 and GFP-RAB13 were seeded on fibronectin-coated glass bottom dishes. At the start of the experiment, the media was exchanged for 200 µl media containing Alexa Fluor 647-conjugated 12G10 antibody (Novus) at a final concentration of 3.7 µg/ml. Samples were fixed with 4% PFA at given time points for subsequent Airyscan imaging at the LSM880 (ZEISS) in the standard super-resolution mode.

### Migration assay

MDA-MB-231 cells were silenced for GGA2 or RAB13 for three days and plated on 25 µg/ml fibronectin-coated 24 well plates. Imaging was performed from 2 hours post plating at the inverted widefield microscope (Nikon Eclipse) using phase-contrast, 10×/0.3 air objective and sCMOS camera (Hamamatsu ORCA-Flash4.0 v3). Single bottom planes of the cells were imaged once every 20 minutes as 2×2 tile for 14 hours. The imaging was performed in full media buffered with 25 mM HEPES pH 7.0–7.6. The plates were kept at 37°C in an atmosphere containing 5% CO_2_. Cell migration was tracked manually in Fiji (Schneider et al., 2012; Schindelin et al., 2012) using the MTrackJ plugin (Meijering et al., 2012). Dividing cells and cells making contacts were not considered for analysis. DiPer Excel macros (Gorelik and Gautreau, 2014) were used to calculate the speed, directionality and to create the visual representation of migration tracks.

### Membrane protrusion speed measurements

MDA-MB-231 cells stably expressing Lifeact-mRFP were created by transduction with lentiviral particles (Jacquemet et al., 2017). pCDH-LifeAct-mRFP plasmid was a gift from P. Caswell (University of Manchester, UK). Lifeact-mRFP cells were silenced for GGA2 or RAB13 for three days and plated on 10 µg/ml fibronectin-coated µ-slide 8 well (Ibidi). Imaging was performed from 2 hours post plating at the Marianas spinning disk confocal microscope (3i) with a CSU-W1 scanning unit (Yokogawa) on an inverted Axio Observer Z1 stage (ZEISS) using a 63×/1.4 Oil Plan-Apochromat objective and Evolve 512 EMCCD camera (Photometrics). The system was controlled by SlideBook 6 software. Single bottom planes of the cells were imaged once every 1 minute for 50–70 minutes. No reduction of cell protrusive activity and photobleaching was observed during the acquisition, suggesting healthy, low phototoxicity imaging conditions. The imaging was performed in full media without phenol red buffered with 25 mM HEPES pH 7.0–7.6. The slides were kept at 37°C in an atmosphere containing 5% CO_2_. The raw images were processed in Fiji using Gaussian blur and cell outline was segmented and membrane speeds calculated in QuimP software version 18.10.01 (Baniukiewicz 2018).

### Inverted invasion assay

200 µl of 5 μg/mL collagen (PureCol® EZ Gel Bovine Collagen Solution, Type I; Advanced Biomatrix, 5074-G) was placed into each transwell chamber (24 Well ThinCert™ Cell Culture Inserts; Greiner Bio-One, 662638) and incubated for 1 h at 37°C in a humidified incubator to set. Transwells were inverted on the lid of a 24-well plate and of 100,000 cells (100 μl volume) in complete medium were placed onto the underside of each transwell filter. Transwells were then covered with the 24-well plate touching the cell suspension drop. The inverted plate was incubated at 37°C for 2–4 h for the cells to attach. Later, transwells were removed, washed twice with serum-free media to remove serum and placed in new 24 well plates containing 1 ml of serum-free media. 100 μl of medium containing 10% FBS as a chemoattractant was pipetted on top of the collagen plug and incubated for 3 days. For visualization, transwells were transferred to a fresh 24-well plate, fixed with 4% PFA for 2 h and stained for actin (phalloidin) after permeabilization with 0.3% (vol/vol) Triton-X 100 for 30 min at room temperature. Virtual 3D Imaging was done with LSM510 confocal microscope (ZEISS) using 20× objective. Individual confocal images were represented in sequence with increasing penetrance of cells from left to right. The invasion was quantified using the area calculator plugin of ImageJ, measuring the fluorescence intensity of cells invading 60?μm or more and expressing this as a percentage of the fluorescence intensity of all cells within the plug.

### Zebrafish xenograft and metastasis assay

Experiments were performed as described in (Pekkonen et al., 2018) under license ESAVI/9339/04.10.07/2016. Briefly, adult casper zebrafishes (roy-/-, mitfa-/-strain) (White et al., 2008) were placed into mating tanks and after natural spawning, the embryos were collected for experiments and cultured in E3-medium (5 mM NaCl, 0.17 mM KCl, 0.33 mM CaCl_2_, 0.33 mM MgSO_4_) supplemented with 0.2 mM phenylthiourea (PTU, Sigma-Aldrich) at 33°C. After two days of incubation, the embryos were anesthetized (MS-222, 200 mg/l, Sigma-Aldrich) and mounted into 0.7% low-melting point agarose for tumor transplantation. Prior to transplantation, siRNA-treated MDA-MB-231 GFP or MDA-MB-231-RFP cells were trypsinized, washed with PBS and resuspended into injection buffer. Approximately 5– 10 nl of cell suspension was microinjected into the pericardial cavity of the embryo using CellTramVario (Eppendorf), Injectman Ni2 (Eppendorf) micromanipulator and glass needles. After transplantation, the embryos were released from the agarose and cultured in E3-PTU at 33°C. On the next day, successfully xenografted healthy embryos were selected and placed into 96-well plates (one embryo per well) and imaged with Nikon Eclipse Ti2 microscope equipped with Plan UW 2× objective. At four days post-injection (dpi) the embryos were anaesthetized and imaged again.

The GFP or RFP (referred as FP) intensity of the tumor was measured manually using FIJI software after background subtraction (rolling ball radius = 25) using Fiji (ImageJ version 1.49 k) (Schneider et al., 2012; Schindelin et al., 2012). The invasion index was calculated as a ratio of the number of invaded cells relative to the FP fluorescence intensity of the primary tumor at 4 dpi. The relative growth of the primary tumor was calculated by dividing FP intensity at 4 dpi by the FP intensity at 1 dpi. Invading cells in the lens were not counted as these sites often autofluoresce. Significantly malformed, dead or poorly oriented embryos were excluded from the analysis. Samples were not blinded for imaging and subsequent analyses.

### In vitro 2D Proliferation assay

Control siRNA or GGA2 siRNA silenced cells were plated on 96-well plates (100 μl; 3000 cells/well). After 1 d, 2 d and 4 d of cell growth, WST-8 reagent (cell counting kit 8, Sigma-Aldrich, 96992) was added to each well (10 μl/well) and absorbance was measured at 450 nm by a plate reader (Thermo, Multiskan Ascent 96/384) after 1 h incubation at +37°C. Culture medium without cells was used to measure background. Relative proliferation was calculated by normalizing the A450 values of 2 d and 4 d to 1 d after background subtraction.

### Mammosphere formation assay

Cells (1000 cells per condition) were plated on 96-well low attachment plates (Corning® Costar® Ultra-Low attachment multiwell plates; Corning, CLS3474-24EA) in Epicult Base medium (StemCell Technologies) supplemented with Epicult supplements, 10 ng/ml EGF, 4 µg/ml heparin and 10 ng/ml bFGF. Cells were allowed to proliferate for 7 days. Mammospheres were counted manually and images captured using the EVOS® Cell Imaging System (ThermoFisher Scientific).

### BioID proximity biotinylation pulldown assay

The protocol was modified from (Roux et al., 2012, 2018) for MDA-MB-231 cells and the BioID-GGA2 construct. Briefly, three 10 cm dishes were transiently transfected per condition (Myc-BioID construct, Myc-BioID-GGA2 construct or mock transfection) in Opti-MEM™ (ThermoFisher, 31985070) containing 50 µM Biotin (Sigma, B4501). After 18 h, media was removed and cells were washed with cold PBS and lysed in 500 µl lysis buffer (0.1% SDS, 0.1% Deoxycholate, 1% TritonX-100, 1 mM EDTA, 200 mM NaCl, 50 mM HEPES pH 7.5, protease and phosphatase inhibitor tablets) for each dish. The lysates were passed through a 27G needle 10 times and incubated for 20 min at 4°C with agitation. Cell lysates were cleared by centrifuge at 10,000×g, 10 min at 4°C and supernatants from the same condition were pooled and added to fresh 2 ml tubes containing magnetic streptavidin beads (Dynabeads® MyOne™ Streptavidin C1; Life Technologies, 65001) pre-washed with 1:3 lysis buffer and incubated overnight at 4°C with agitation. On the following day, supernatants were removed from the tubes and beads were washed once in buffer1 (0.5% deoxycholate, 0.5% NP-40, 1 mM EDTA, 250 mM LiCl, 10 mM Tris-Cl pH 7.4, protease and phosphatase inhibitor tablets) for 5 min and then in buffer2 (200 mM NaCl, 75 mM Tris-Cl pH 7.4, 7.5 mM EDTA, 7.5 mM EGTA, 1.5% TritonX-100, 0.75% NP-40, protease and phosphatase inhibitor tablets) for 5 min. Finally, proteins were eluted from the beads in 50 µl of reducing SDS sample buffer for 5 min at 95°C and separated by SDS-PAGE. The gel was then stained with Coomassie blue and each lane was cut carefully into 5–8 pieces for in gel-digestion and mass spectrometry analysis.

### Mass spectrometry analysis

Proteins cut from an SDS-PAGE gel were in-gel digested by trypsin according to a standard protocol (Shevchenko et al., 1996). The LC-ESI-MS/MS analyses were performed on a nanoflow HPLC system (Easy-nLCII, Thermo Fisher Scientific) coupled to the LTQ Orbitrap Velos Pro mass spectrometer (Thermo Fisher Scientific, Bremen, Germany) equipped with a nano-electrospray ionization source. Peptides were first loaded on a trapping column and subsequently separated inline on a 15 cm C18 column (75 μm x 15 cm, ReproSil-Pur 5 μm 200 Å C18-AQ, Dr. Maisch HPLC GmbH, Ammerbuch-Entringen, Germany). The mobile phase consisted of water/acetonitrile (98:2 (v/v)) with 0.1% formic acid (solvent A) and acetonitrile/water (95:5 (v/v)) with 0.1% formic acid (solvent B). A two-step 45 min gradient from 5% to 40% B was used to elute peptides.

MS data were acquired automatically by using Thermo Xcalibur 3.0 software (Thermo Fisher Scientific). An information dependent acquisition method consisted of an Orbitrap MS survey scan of mass range 300-2000 m/z followed by CID fragmentation for 10 most intense peptide ions. Fragmentation, and detection of fragment ions (MS2 scan), was performed in a linear ion trap.

The data files were searched for protein identification using Proteome Discoverer 1.4 software (Thermo Fisher Scientific) connected to an in-house Mascot server running the Mascot 2.5 software (Matrix Science). Data were searched against the SwissProt database (release 2015_08). The search included carbamidomethyl as a fixed modification and methionine oxidation as variable modification together with one tryptic missed cleavage. The peptide and protein false discovery rate (FDR) was set to 5%. Proteins assigned at least with two peptides were accepted.

### Genomic Analyses of GGA2 and RAB13 from the Public Datasets

For investigating the differential expression pattern of GGA2 and RAB13 mRNA, TCGA tumor samples were compared with paired adjacent TCGA and GTEx normal samples using the GEPIA2 platform (http://gepia2.cancer-pku.cn/). GEPIA2 is an updated version of GEPIA (Tang et al., 2017) for analyzing the RNA sequencing expression data of 9,736 tumors and 8,587 normal samples from the TCGA and the GTEx projects, using a standard processing pipeline. The expression data are first log_2_(TPM+1) transformed for differential analysis and the log_2_ fold change (cut-off > 0.25) is defined as median(Tumor)-median(Normal).

### Online supplemental material

Fig. S1 shows the results of the siRNA screen identifying a role for GGA2 in integrin trafficking in addition to control endocytosis experiments. Fig. S2 shows the localization of fluorescent protein fusions of GGA2 in comparison to endosomal markers. Fig. S3 shows supporting information for the BioID experiment. Fig. S4 shows integrin trafficking experiments in response to RAB10 and SEC23B knockdown and the effect of RAB13 on inactive β1-integrin traffic. Video 1 shows random 2D migration of control and GGA2 or RAB13 depleted cells. Video 2 shows lamellipodial dynamics of control and GGA2 or RAB13 depleted cells. Additionally, a table summarizing all used reagents and resources is available as a MS Word document and a table of mass spectrometry hits for GGA2 interactors is available as an excel file.

## Supplementary figures 1-4 and associated figure legends

**Supplementary Figure 1.**
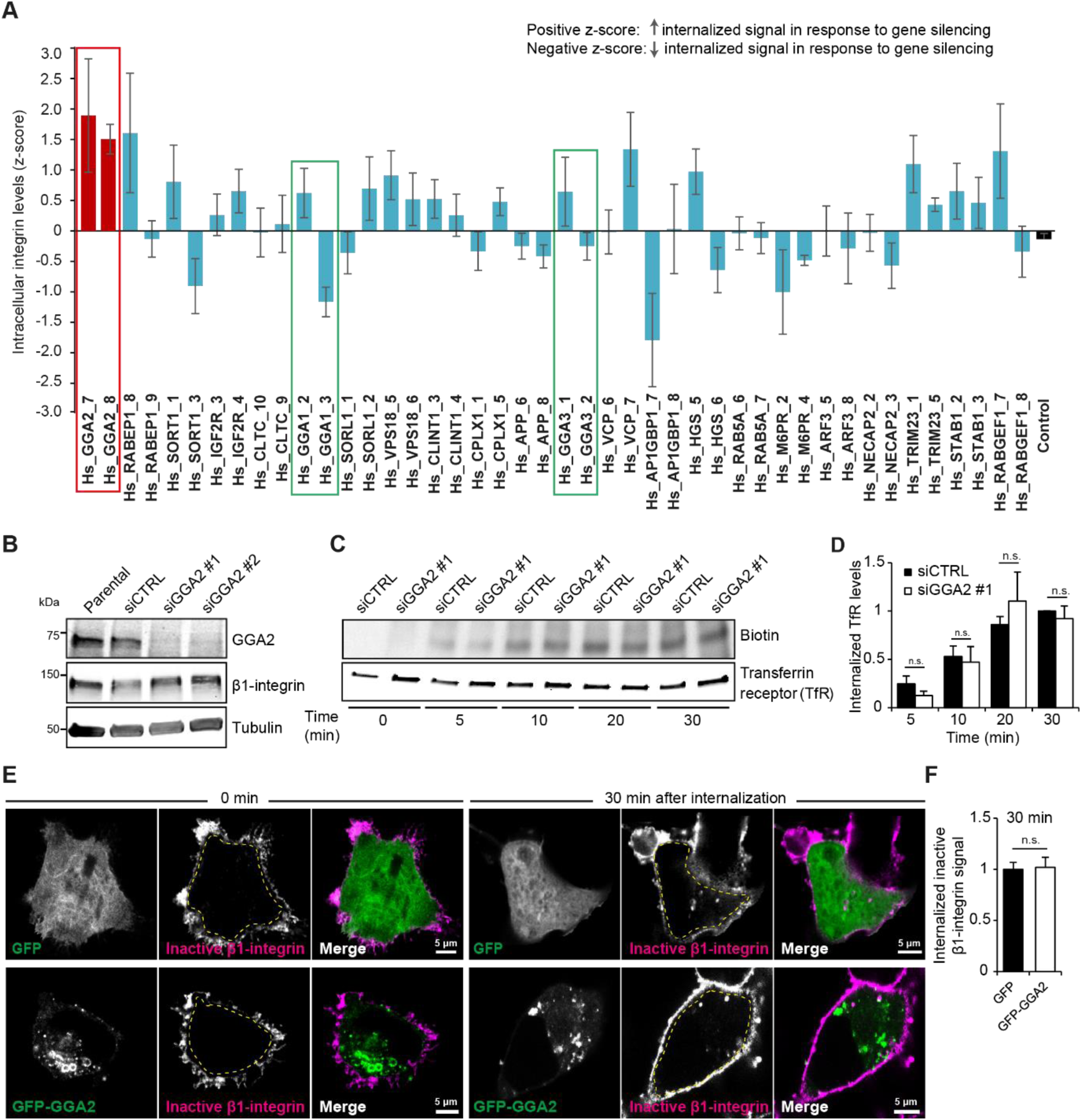
RNAi-based screening identifies GGA2 as a regulator of integrin internalization. **A)** Differential effects of human GGAs and their known interactors on β1-integrin internalization. After silencing of the indicated genes with two siRNAs using the CSMA platform (see methods), β1-integrin endocytosis was measured using the antibody-based β1-integrin endocytosis assay at 30 min internalization. Endocytosis levels are represented as Z-scores for the whole dataset (red bars: GGA2 siRNAs; green outlines: GGA1 and GGA3 siRNAs; cyan bars: siRNAs against GGA interactors; black bar: control siRNA). Positive z-scores indicate increased internalization of β1-integrin in response to gene silencing. (Data are mean ± SEM; n = 4). **B)** Representative western blot analysis of GGA2 and β1-integrin levels in non-transfected parental cells and in siCTRL, siGGA2 #1 and siGGA2 #2 transfected MDA-MB-231 cells. Tubulin is included as a loading control. **C-D)** Internalization of biotinylated cell-surface transferrin receptor (TfR) in siCTRL or siGGA2 #1 MDA-MB-231 cells. Shown is a representative western blot and quantification of biotinylated TfR relative to total TfR normalized to siCTRL at 30 min time point (data are mean ± SEM of n = 3 independent experiments; statistical analysis, Student’s unpaired *t*-test; n.s. = not significant) **E-F)** Representative confocal images (E) and quantification (F, performed as in Fig. 1L) of internalized inactive β1-integrin (MAB13 antibody) at the indicated time points in MDA-MB-231 cells expressing GFP and GFP-GGA2 (data are mean ± SEM; n = 70-110 cells per condition). Statistical analysis, Student’s unpaired *t*-test (n.s.= not significant).

**Supplementary Figure 2.**
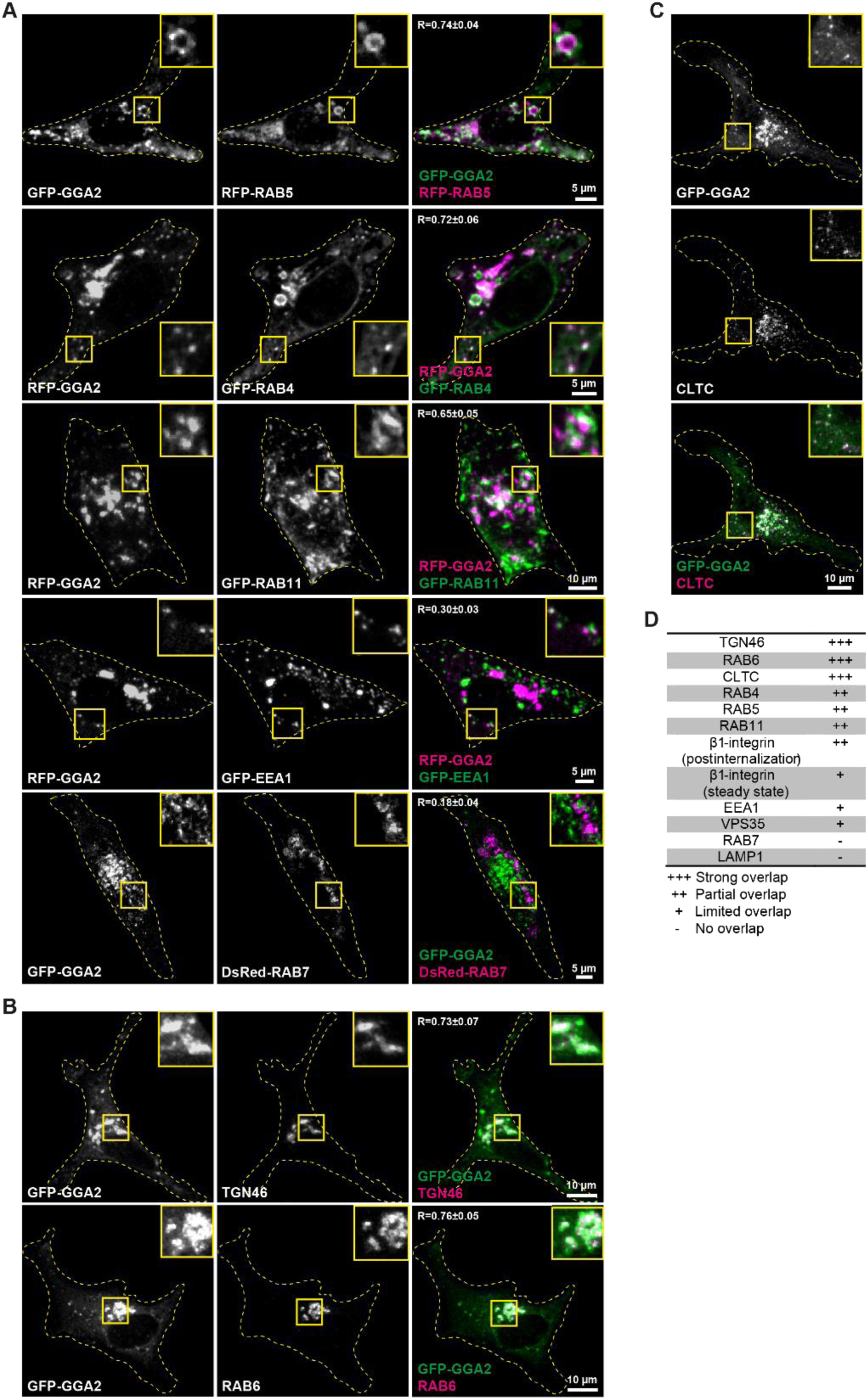
Cellular localization of fluorescently tagged GGA2. **A)** Representative confocal microscopy images of MDA-MB-231 cells co-transfected with the indicated fluorescently tagged vesicular compartment proteins (proteins of interest, magenta) and fluorescently tagged GGA2. Overlap between the indicated proteins is quantified (R ± SD; n = 7 - 10 cells per condition). **B-C)** Representative confocal microscopy images of MDA-MB-231 cells transfected with GFP-GGA2 and stained for intracellular compartments with reported GGA2 localization. Overlap between the indicated proteins is quantified (R ± SD; n = 7 - 10 cells per condition). TGN46: Trans-Golgi network integral membrane protein 2; CLTC: clathrin heavy chain. **D)** Analysis of GGA2 (endogenous or fluorescently tagged) overlap with different endogenous markers (vesicles or organelles) at steady state based on R-values and visual comparison (images analyzed from Fig 3 and Fig S2).

**Supplementary Figure 3.**
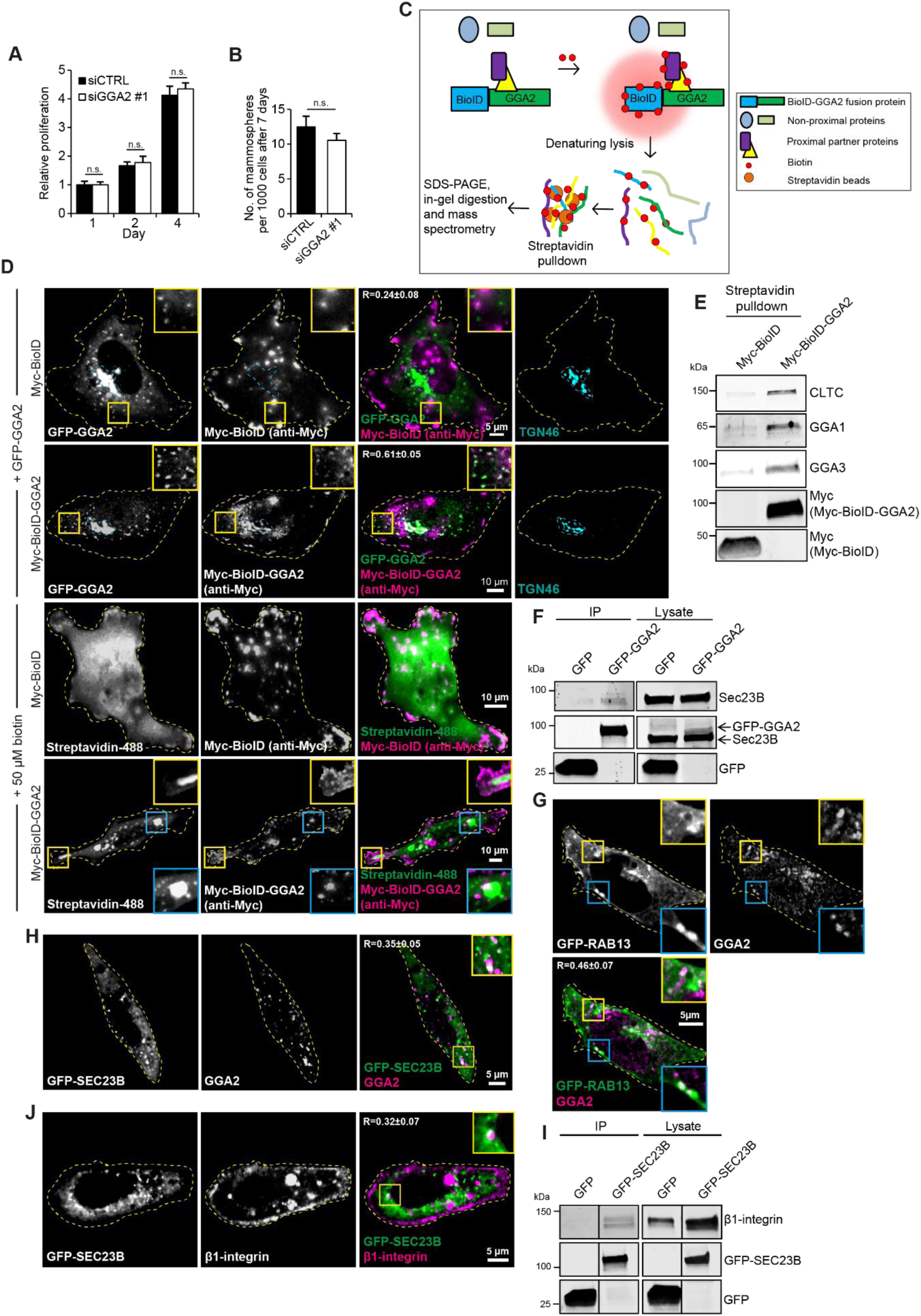
Validation of the myc-BioID-GGA2 fusion protein. **A)** Quantification of siCTRL or siGGA2 #1 MDA-MB-231 cell proliferation over 4 days (data are mean ± SEM; n = 3 independent experiments). **B)** Quantification of the number of tumor spheres formed by siCTRL or siGGA2 #1 MDA-MB-231 cells (data are mean ± SEM; n = 3 independent experiments). **C)** Outline of the BioID approach to detect new GGA2-interacting proteins. **D)** Representative confocal microscopy images of MDA-MB-231 cells expressing Myc-BioID or Myc-BioID-GGA2 co-transfected with GFP-GGA2 or transfected in the presence of 50 µM biotin. TGN46 was used as positive control for correct Myc-BioID-GGA2 localization. Addition of biotin (signal detected with Alexa Fluor 488-conjugated streptavidin) was used to demonstrate that biotinylation occurs in the same cellular compartments as Myc-BioID-GGA2 localization. Overlap between the indicated proteins is quantified (R ± SD) and highlighted in both yellow and blue insets. **E)** Streptavidin pulldown in MDA-MB-231 cells transiently transfected with Myc-BioID or Myc-BioID-GGA2, treated with biotin and lysed. Immunoblotting for known GGA2 interactors (CLTC, GGA1, GGA3) and Myc tag is shown. **F)** Representative western blot of GFP-Trap pulldown in MDA-MB-231 cells transiently transfected with GFP or GFP-GGA2 and analyzed for endogenous SEC23B and for GFP (n = 3). Membranes in the lysate were first probed with an antibody against SEC23B and then for GFP. SEC23B and GFP-GGA2 are of a similar size and thus appear at the same level in the membrane and are indicated by arrows. **G)** Representative confocal image of an MDA-MB-231 cell transiently expressing GFP-RAB13 and stained with anti-GGA2 antibody. Overlap between the indicated proteins is highlight in yellow and blue insets (n = 80-200 cells per condition). **H)** Representative confocal images of MDA-MB-231 cells expressing GFP-SEC23B and stained with anti-GGA2 antibody. Overlap between the indicated proteins is highlighted in the yellow inset (n = 80-200 cells per condition). **I)** Representative western blot of GFP-Trap pulldown in MDA-MB-231 cells transiently expressing GFP-SEC23B and analyzed for endogenous β1-integrin and for GFP (n = 3). **J)** Representative confocal microscopy images of MDA-MB-231 cells transfected with GFP-SEC23B and stained with β1-integrin antibody P5D2. Overlap between the indicated proteins is highlighted in the yellow inset and quantified (R ± SD; n = 150 cells).

**Supplementary Figure 4.**
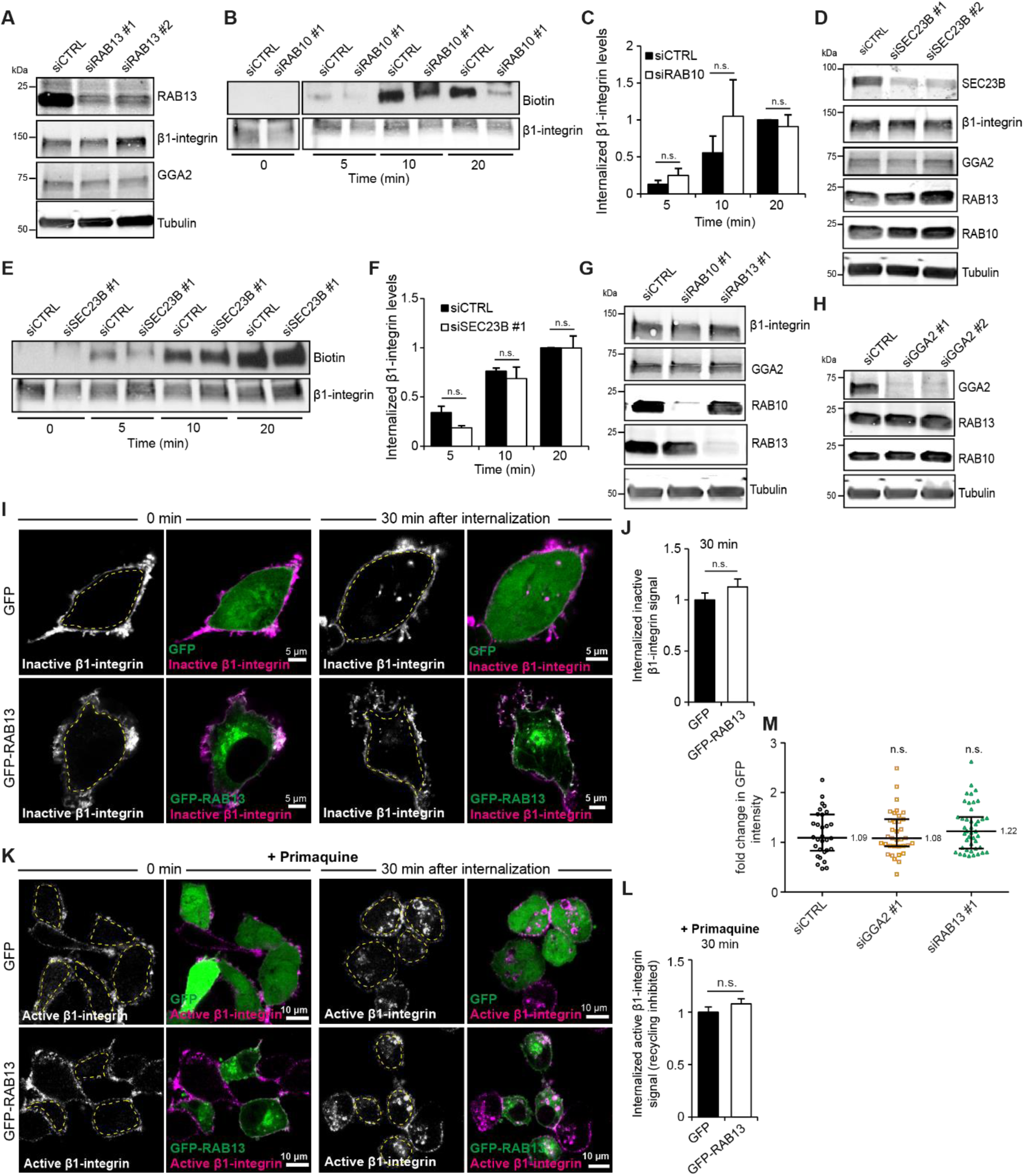
RAB10 and SEC23B do not regulate β1-integrin traffic and RAB13 function is specific for active β1-integrin recycling. **A)** Representative western blots of MDA-MB-231 cells silenced with control siRNA, RAB13 siRNA#1 or RAB13 siRNA#2 and blotted for β1-integrin and GGA2. Tubulin was used as a loading control. **B-C)** Internalization of biotinylated cell-surface β1-integrin in control siRNA or RAB10 siRNA silenced MDA-MB-231 cells. Shown is a representative western blot and quantification of biotinylated β1-integrin relative to total β1-integrin, normalized against 20 min siCTRL time point (data are mean ± SEM of n = 3 independent experiments). **D)** Representative western blot of β1-integrin, GGA2, RAB13 and RAB10 levels in MDA-MB-231 cells silenced with control siRNA, SEC23B siRNA#1 or SEC23B siRNA#2. Tubulin was used as a loading control. **E-F)** Internalization of biotinylated cell-surface β1-integrin in MDA-MB-231 cells treated with either control siRNA (siCTRL) or SEC23B siRNA oligo #1 (siSEC23B #1). Shown is a representative western blot (E) and quantification of biotinylated β1-integrin relative to total β1-integrin levels and normalized to siCTRL 20 min time point (F, data are mean ± S.E.M.; n = 3) **G)** Representative western blots of MDA-MB-231 cells silenced with control siRNA, RAB10 siRNA#1 or RAB13 siRNA#1 and analyzed as indicated. Tubulin was used as a loading control. **H)** Representative western blots of MDA-MB-231 cells silenced with control siRNA, GGA2 siRNA#1 or GGA2 siRNA#2 and analyzed as indicated. Tubulin was used as a loading control. **I-J)** Representative confocal microscopy images (A) and quantification (B, performed as in Fig. 1L) of internalized inactive β1-integrin (MAB13 antibody) in MDA-MB-231 cells transfected with either GFP or GFP-RAB13 (data are mean ± SEM.; n = 55-110 cells per condition). **K-L)** Representative confocal microscopy images (K) and quantification (L) of internalized active β1-integrin (12G10 antibody) in MDA-MB-231 cells transfected with either GFP or GFP-RAB13 and treated with 100 μm primaquine (data are mean ± SEM.; n = 70-80 cells per condition). Statistical analysis for quantifications (n.s.= not significant, **p < 0.005, student’s unpaired *t*-test).

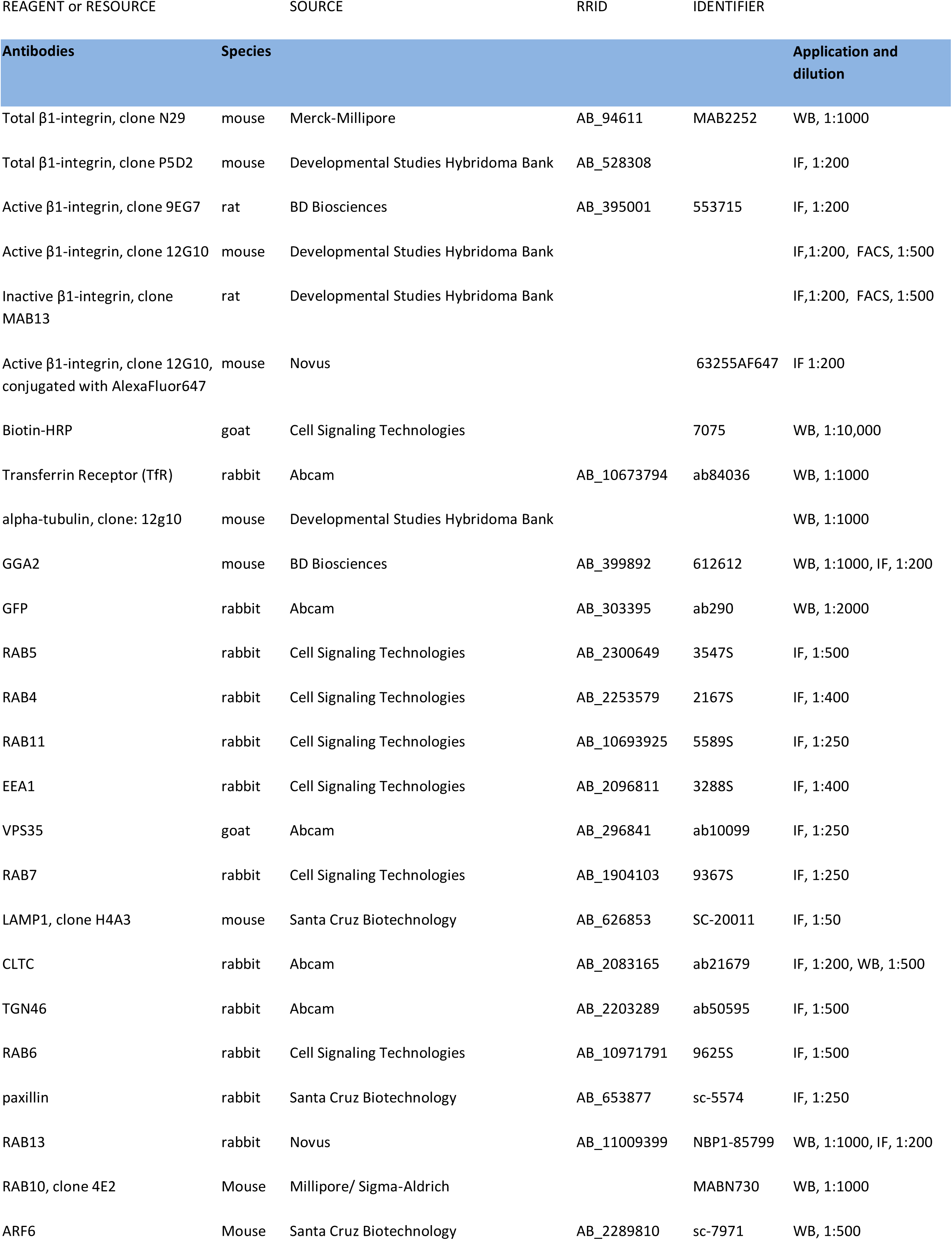
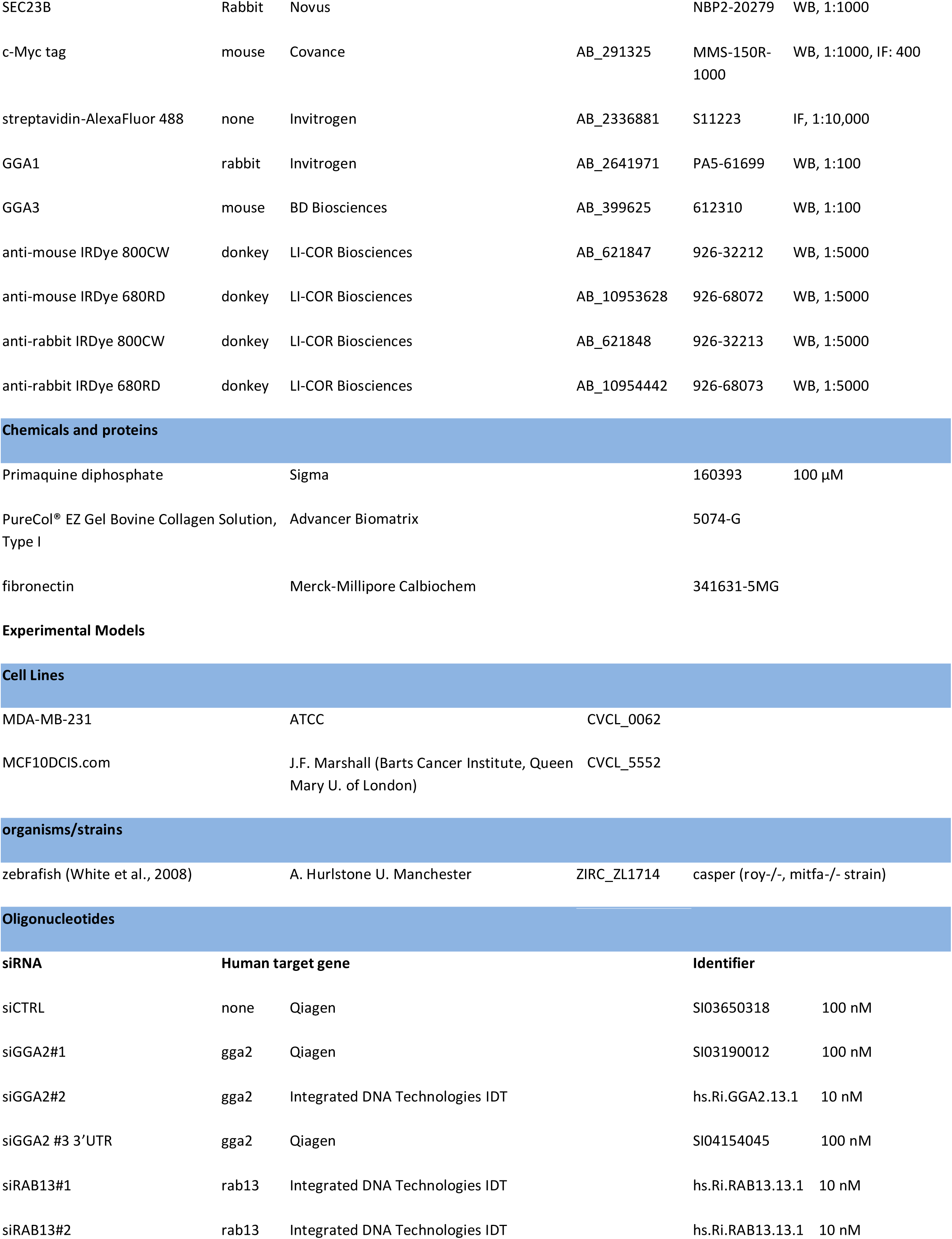
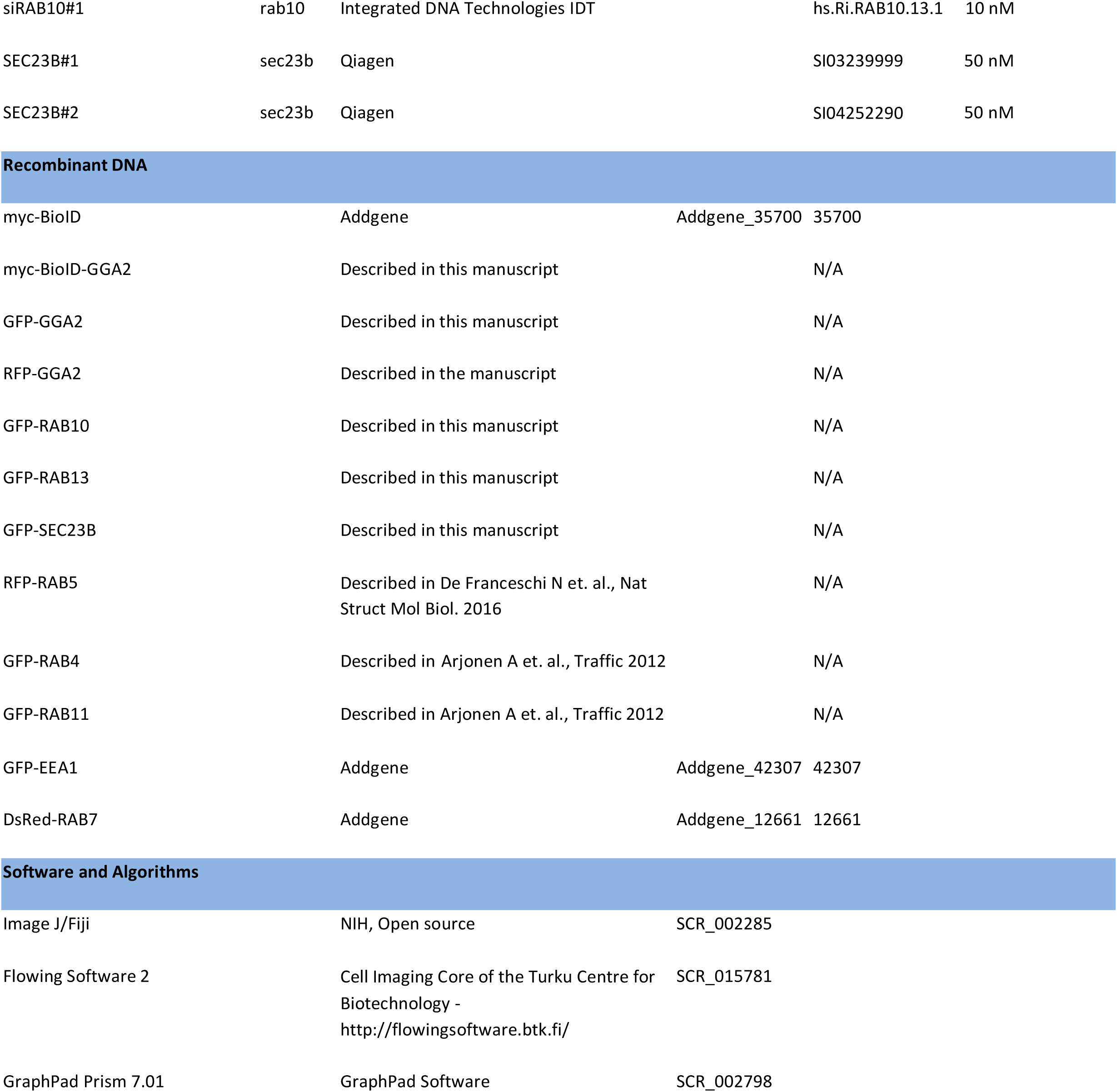

## References

Alanko, J., A. Mai, G. Jacquemet, K. Schauer, R. Kaukonen, M. Saari, B. Goud, and J. Ivaska. 2015. Integrin endosomal signalling suppresses anoikis. Nat. Cell Biol. 17:1412–1421. doi:10.1038/ncb3250.

Argenzio, E., C. Margadant, D. Leyton-Puig, H. Janssen, K. Jalink, A. Sonnenberg, and W.H. Moolenaar. 2014. CLIC4 regulates cell adhesion and β1 integrin trafficking. J. Cell Sci. 127:5189–5203. doi:10.1242/jcs.150623.

Arjonen, A., J. Alanko, S. Veltel, and J. Ivaska. 2012. Distinct recycling of active and inactive β1 integrins. Traffic. 13:610–625. doi:10.1111/j.1600-0854.2012.01327.x.

Barlowe, C., L. Orci, T. Yeung, M. Hosobuchi, S. Hamamoto, N. Salama, M.F. Rexach, M. Ravazzola, M. Amherdt, and R. Schekman. 1994. COPII: a membrane coat formed by Sec proteins that drive vesicle budding from the endoplasmic reticulum. Cell. 77:895–907.

Boman, A.L., C. j Zhang, X. Zhu, and R.A. Kahn. 2000. A family of ADP-ribosylation factor effectors that can alter membrane transport through the trans-Golgi. Mol. Biol. Cell. 11:1241–1255. doi:10.1091/mbc.11.4.1241.

Böttcher, R.T., C. Stremmel, A. Meves, H. Meyer, M. Widmaier, H.-Y. Tseng, and R. Fässler. 2012. Sorting nexin 17 prevents lysosomal degradation of β1 integrins by binding to the β1-integrin tail. Nat. Cell Biol. 14:584–592. doi:10.1038/ncb2501.

Bridgewater, R.E., J.C. Norman, and P.T. Caswell. 2012. Integrin trafficking at a glance. J Cell Sci. 125:3695–3701. doi:10.1242/jcs.095810.

Bruno, J., A. Brumfield, N. Chaudhary, D. Iaea, and T.E. McGraw. 2016. SEC16A is a RAB10 effector required for insulin-stimulated GLUT4 trafficking in adipocytes. J. Cell Biol. 214:61–76. doi:10.1083/jcb.201509052.

Calderwood, D.A., I.D. Campbell, and D.R. Critchley. 2013. Talins and kindlins: partners in integrin-mediated adhesion. Nat. Rev. Mol. Cell Biol. 14:503–517. doi:10.1038/nrm3624.

Campbell, I.D., and M.J. Humphries. 2011. Integrin Structure, Activation, and Interactions. Cold Spring Harb Perspect Biol. 3:a004994. doi:10.1101/cshperspect.a004994.

Caswell, P.T., M. Chan, A.J. Lindsay, M.W. McCaffrey, D. Boettiger, and J.C. Norman. 2008. Rab-coupling protein coordinates recycling of α5β1 integrin and EGFR1 to promote cell migration in 3D microenvironments. J Cell Biol. 183:143–155. doi:10.1083/jcb.200804140.

Caswell, P.T., H.J. Spence, M. Parsons, D.P. White, K. Clark, K.W. Cheng, G.B. Mills, M.J. Humphries, A.J. Messent, K.I. Anderson, M.W. McCaffrey, B.W. Ozanne, and J.C. Norman. 2007. Rab25 Associates with α5β1 Integrin to Promote Invasive Migration in 3D Microenvironments. Dev. Cell. 13:496–510. doi:10.1016/j.devcel.2007.08.012.

Chen, P.-W., R. Luo, X. Jian, and P.A. Randazzo. 2014. The Arf6 GTPase-activating proteins ARAP2 and ACAP1 define distinct endosomal compartments that regulate integrin α5β1 traffic. J. Biol. Chem. 289:30237–30248. doi:10.1074/jbc.M114.596155.

De Franceschi, N., A. Arjonen, N. Elkhatib, K. Denessiouk, A.G. Wrobel, T.A. Wilson, J. Pouwels, G. Montagnac, D.J. Owen, and J. Ivaska. 2016. Selective integrin endocytosis is driven by interactions between the integrin α-chain and AP2. Nat. Struct. Mol. Biol. 23:172–179. doi:10.1038/nsmb.3161.

De Franceschi, N., H. Hamidi, J. Alanko, P. Sahgal, and J. Ivaska. 2015. Integrin traffic-the update. J. Cell Sci. 128:839–852. doi:10.1242/jcs.161653.

Dell’Angelica, E.C., R. Puertollano, C. Mullins, R.C. Aguilar, J.D. Vargas, L.M. Hartnell, and J.S. Bonifacino. 2000. GGAs: a family of ADP ribosylation factor-binding proteins related to adaptors and associated with the Golgi complex. J. Cell Biol. 149:81–94.

Diggins, N.L., H. Kang, A. Weaver, and D.J. Webb. 2018. α5β1 integrin trafficking and Rac activation are regulated by APPL1 in a Rab5-dependent manner to inhibit cell migration. J. Cell Sci. 131. doi:10.1242/jcs.207019.

Dozynkiewicz, M.A., N.B. Jamieson, I. Macpherson, J. Grindlay, P.V.E. van den Berghe, A. von Thun, J.P. Morton, C. Gourley, P. Timpson, C. Nixon, C.J. McKay, R. Carter, D. Strachan, K. Anderson, O.J. Sansom, P.T. Caswell, and J.C. Norman. 2012. Rab25 and CLIC3 collaborate to promote integrin recycling from late endosomes/lysosomes and drive cancer progression. Dev. Cell. 22:131–145. doi:10.1016/j.devcel.2011.11.008.

Dunphy, J.L., R. Moravec, K. Ly, T.K. Lasell, P. Melancon, and J.E. Casanova. 2006. The Arf6 GEF GEP100/BRAG2 regulates cell adhesion by controlling endocytosis of beta1 integrins. Curr. Biol. 16:315–320. doi:10.1016/j.cub.2005.12.032.

Fang, Z., N. Takizawa, K.A. Wilson, T.C. Smith, A. Delprato, M.W. Davidson, D.G. Lambright, and E.J. Luna. 2010. The membrane-associated protein, supervillin, accelerates F-actin-dependent rapid integrin recycling and cell motility. Traffic. 11:782–799. doi:10.1111/j.1600-0854.2010.01062.x.

Felding-Habermann, B., T.E. O’Toole, J.W. Smith, E. Fransvea, Z.M. Ruggeri, M.H. Ginsberg, P.E. Hughes, N. Pampori, S.J. Shattil, A. Saven, and B.M. Mueller. 2001. Integrin activation controls metastasis in human breast cancer. Proc. Natl. Acad. Sci. U.S.A. 98:1853–1858. doi:10.1073/pnas.98.4.1853.

Fletcher, S.J., and J.Z. Rappoport. 2010. Moving forward: polarised trafficking in cell migration. Trends Cell Biol. 20:71–78. doi:10.1016/j.tcb.2009.11.006.

Ghosh, P., J. Griffith, H.J. Geuze, and S. Kornfeld. 2003. Mammalian GGAs act together to sort mannose 6-phosphate receptors. J. Cell Biol. 163:755–766. doi:10.1083/jcb.200308038.

Gorelick, F.S., and C. Shugrue. 2001. Exiting the endoplasmic reticulum. Mol. Cell. Endocrinol. 177:13–18.

Gorelik, R., and A. Gautreau. 2014. Quantitative and unbiased analysis of directional persistence in cell migration. Nat Protoc. 9:1931–1943. doi:10.1038/nprot.2014.131.

Govero, J., B. Doray, H. Bai, and S. Kornfeld. 2012. Analysis of Gga Null Mice Demonstrates a Non-Redundant Role for Mammalian GGA2 during Development. PLOS ONE. 7:e30184. doi:10.1371/journal.pone.0030184.

Gu, Z., E.H. Noss, V.W. Hsu, and M.B. Brenner. 2011. Integrins traffic rapidly via circular dorsal ruffles and macropinocytosis during stimulated cell migration. J. Cell Biol. 193:61–70. doi:10.1083/jcb.201007003.

Hegde, S., and S. Raghavan. 2013. A skin-depth analysis of integrins: role of the integrin network in health and disease. Cell Commun. Adhes. 20:155–169. doi:10.3109/15419061.2013.854334.

Hirst, J., G.H.H. Borner, R. Antrobus, A.A. Peden, N.A. Hodson, D.A. Sahlender, and M.S. Robinson. 2012. Distinct and overlapping roles for AP-1 and GGAs revealed by the “knocksideways” system. Curr. Biol. 22:1711–1716. doi:10.1016/j.cub.2012.07.012.

Hirst, J., W.W. Lui, N.A. Bright, N. Totty, M.N. Seaman, and M.S. Robinson. 2000. A family of proteins with gamma-adaptin and VHS domains that facilitate trafficking between the trans-Golgi network and the vacuole/lysosome. J. Cell Biol. 149:67–80.

Hynes, R.O. 2002. Integrins: bidirectional, allosteric signaling machines. Cell. 110:673–687.

Ioannou, M.S., E.S. Bell, M. Girard, M. Chaineau, J.N.R. Hamlin, M. Daubaras, A. Monast, M. Park, L. Hodgson, and P.S. McPherson. 2015. DENND2B activates Rab13 at the leading edge of migrating cells and promotes metastatic behavior. J. Cell Biol. 208:629–648. doi:10.1083/jcb.201407068.

Jacquemet, G., I. Paatero, A.F. Carisey, A. Padzik, J.S. Orange, H. Hamidi, and J. Ivaska. 2017. FiloQuant reveals increased filopodia density during breast cancer progression. J. Cell Biol. 216:3387–3403. doi:10.1083/jcb.201704045.

Jin, J.-K., P.-C. Tien, C.-J. Cheng, J.H. Song, C. Huang, S.-H. Lin, and G.E. Gallick. 2015. Talin1 phosphorylation activates β1 integrins: a novel mechanism to promote prostate cancer bone metastasis. Oncogene. 34:1811–1821. doi:10.1038/onc.2014.116.

Jonker, C.T.H., R. Galmes, T. Veenendaal, C. Ten Brink, R.E.N. van der Welle, N. Liv, J. de Rooij, A.A. Peden, P. van der Sluijs, C. Margadant, and J. Klumperman. 2018. Vps3 and Vps8 control integrin trafficking from early to recycling endosomes and regulate integrin-dependent functions. Nat. Commun. 9:792. doi:10.1038/s41467-018-03226-8.

Kawauchi, T. 2012. Cell adhesion and its endocytic regulation in cell migration during neural development and cancer metastasis. Int J Mol Sci. 13:4564–4590. doi:10.3390/ijms13044564.

Liu, J., M. Das, J. Yang, S.S. Ithychanda, V.P. Yakubenko, E.F. Plow, and J. Qin. 2015. Structural mechanism of integrin inactivation by filamin. Nat. Struct. Mol. Biol. 22:383–389. doi:10.1038/nsmb.2999.

Mai, A., S. Veltel, T. Pellinen, A. Padzik, E. Coffey, V. Marjomäki, and J. Ivaska. 2011. Competitive binding of Rab21 and p120RasGAP to integrins regulates receptor traffic and migration. J. Cell Biol. 194:291–306. doi:10.1083/jcb.201012126.

McNally, K.E., R. Faulkner, F. Steinberg, M. Gallon, R. Ghai, D. Pim, P. Langton, N. Pearson, C.M. Danson, H. Nägele, L.L. Morris, A. Singla, B.L. Overlee, K.J. Heesom, R. Sessions, L. Banks, B.M. Collins, I. Berger, D.D. Billadeau, E. Burstein, and P.J. Cullen. 2017. Retriever is a multiprotein complex for retromer-independent endosomal cargo recycling. Nat. Cell Biol. 19:1214–1225. doi:10.1038/ncb3610.

Meijering, E., O. Dzyubachyk, and I. Smal. 2012. Methods for cell and particle tracking. Meth. Enzymol. 504:183–200. doi:10.1016/B978-0-12-391857-4.00009-4.

Moreno-Layseca, P., J. Icha, H. Hamidi, and J. Ivaska. 2019. Integrin trafficking in cells and tissues. Nat. Cell Biol. 21:122–132. doi:10.1038/s41556-018-0223-z.

Morgan, M.R., H. Hamidi, M.D. Bass, S. Warwood, C. Ballestrem, and M.J. Humphries. 2013. Syndecan-4 phosphorylation is a control point for integrin recycling. Dev. Cell. 24:472–485. doi:10.1016/j.devcel.2013.01.027.

Nader, G.P.F., E.J. Ezratty, and G.G. Gundersen. 2016. FAK, talin and PIPKI? regulate endocytosed integrin activation to polarize focal adhesion assembly. Nature Cell Biology. 18:491–503. doi:10.1038/ncb3333.

Nielsen, M.S., P. Madsen, E.I. Christensen, A. Nykjaer, J. Gliemann, D. Kasper, R. Pohlmann, and C.M. Petersen. 2001. The sortilin cytoplasmic tail conveys Golgi-endosome transport and binds the VHS domain of the GGA2 sorting protein. EMBO J. 20:2180–2190. doi:10.1093/emboj/20.9.2180.

Nishikimi, A., S. Ishihara, M. Ozawa, K. Etoh, M. Fukuda, T. Kinashi, and K. Katagiri. 2014. Rab13 acts downstream of the kinase Mst1 to deliver the integrin LFA-1 to the cell surface for lymphocyte trafficking. Sci. Signal. 7:ra72. doi:10.1126/scisignal.2005199.

Nokes, R.L., I.C. Fields, R.N. Collins, and H. Fölsch. 2008. Rab13 regulates membrane trafficking between TGN and recycling endosomes in polarized epithelial cells. J. Cell Biol. 182:845–853. doi:10.1083/jcb.200802176.

O’Farrell, H., B. Harbourne, Z. Kurlawala, Y. Inoue, A.L. Nagelberg, V.D. Martinez, D. Lu, M.H. Oh, B.P. Coe, K.L. Thu, R. Somwar, S. Lam, W.L. Lam, A.M. Unni, L. Beverly, and W.W. Lockwood. 2018. Integrative Genomic Analyses Identifies GGA2 as a Cooperative Driver of EGFR-Mediated Lung Tumorigenesis. J. Thorac. Oncol. doi:10.1016/j.jtho.2018.12.004.

Parachoniak, C.A., Y. Luo, J.V. Abella, J.H. Keen, and M. Park. 2011. GGA3 functions as a switch to promote Met receptor recycling, essential for sustained ERK and cell migration. Dev. Cell. 20:751–763. doi:10.1016/j.devcel.2011.05.007.

Parachoniak, C.A., and M. Park. 2012. Dynamics of receptor trafficking in tumorigenicity. Trends Cell Biol. 22:231–240. doi:10.1016/j.tcb.2012.02.002.

Paul, N.R., J.L. Allen, A. Chapman, M. Morlan-Mairal, E. Zindy, G. Jacquemet, L. Fernandez del Ama, N. Ferizovic, D.M. Green, J.D. Howe, E. Ehler, A. Hurlstone, and P.T. Caswell. 2015a. α5β1 integrin recycling promotes Arp2/3-independent cancer cell invasion via the formin FHOD3. J. Cell Biol. 210:1013–1031. doi:10.1083/jcb.201502040.

Paul, N.R., G. Jacquemet, and P.T. Caswell. 2015b. Endocytic Trafficking of Integrins in Cell Migration. Curr. Biol. 25:R1092-1105. doi:10.1016/j.cub.2015.09.049.

Pekkonen, P., S. Alve, G. Balistreri, S. Gramolelli, O. Tatti-Bugaeva, I. Paatero, O. Niiranen, K. Tuohinto, N. Perälä, A. Taiwo, N. Zinovkina, P. Repo, K. Icay, J. Ivaska, P. Saharinen, S. Hautaniemi, K. Lehti, and P.M. Ojala. 2018. Lymphatic endothelium stimulates melanoma metastasis and invasion via MMP14-dependent Notch3 and β1-integrin activation. Elife. 7. doi:10.7554/eLife.32490.

Pellinen, T., A. Arjonen, K. Vuoriluoto, K. Kallio, J.A.M. Fransen, and J. Ivaska. 2006. Small GTPase Rab21 regulates cell adhesion and controls endosomal traffic of beta1-integrins. J. Cell Biol. 173:767–780. doi:10.1083/jcb.200509019.

Powelka, A.M., J. Sun, J. Li, M. Gao, L.M. Shaw, A. Sonnenberg, and V.W. Hsu. 2004. Stimulation-Dependent Recycling of Integrin β1 Regulated by ARF6 and Rab11. Traffic. 5:20–36. doi:10.1111/j.1600-0854.2004.00150.x.

Pozzi, A., and R. Zent. 2013. Integrins in kidney disease. J. Am. Soc. Nephrol. 24:1034–1039. doi:10.1681/ASN.2013010012.

Puertollano, R., and J.S. Bonifacino. 2004. Interactions of GGA3 with the ubiquitin sorting machinery. Nat. Cell Biol. 6:244–251. doi:10.1038/ncb1106.

Puertollano, R., P.A. Randazzo, J.F. Presley, L.M. Hartnell, and J.S. Bonifacino. 2001. The GGAs promote ARF-dependent recruitment of clathrin to the TGN. Cell. 105:93–102.

Rainero, E., and J.C. Norman. 2015. Endosomal integrin signals for survival. Nat. Cell Biol. 17:1373–1375. doi:10.1038/ncb3261.

Rantala, J.K., R. Mäkelä, A.-R. Aaltola, P. Laasola, J.-P. Mpindi, M. Nees, P. Saviranta, and O. Kallioniemi. 2011a. A cell spot microarray method for production of high density siRNA transfection microarrays. BMC Genomics. 12:162. doi:10.1186/1471-2164-12-162.

Rantala, J.K., J. Pouwels, T. Pellinen, S. Veltel, P. Laasola, E. Mattila, C.S. Potter, T. Duffy, J.P. Sundberg, O. Kallioniemi, J.A. Askari, M.J. Humphries, M. Parsons, M. Salmi, and J. Ivaska. 2011b. SHARPIN is an endogenous inhibitor of β1-integrin activation. Nat. Cell Biol. 13:1315–1324. doi:10.1038/ncb2340.

Ratcliffe, C.D.H., P. Sahgal, C.A. Parachoniak, J. Ivaska, and M. Park. 2016. Regulation of Cell Migration and β1 Integrin Trafficking by the Endosomal Adaptor GGA3. Traffic. 17:670–688. doi:10.1111/tra.12390.

Roux, K.J., D.I. Kim, B. Burke, and D.G. May. 2018. BioID: A Screen for Protein-Protein Interactions. Curr. Protoc. Protein Sci. 91:19.23.1-19.23.15. doi:10.1002/cpps.51.

Roux, K.J., D.I. Kim, M. Raida, and B. Burke. 2012. A promiscuous biotin ligase fusion protein identifies proximal and interacting proteins in mammalian cells. J. Cell Biol. 196:801–810. doi:10.1083/jcb.201112098.

Sakurai, A., J. Gavard, Y. Annas-Linhares, J.R. Basile, P. Amornphimoltham, T.R. Palmby, H. Yagi, F. Zhang, P.A. Randazzo, X. Li, R. Weigert, and J.S. Gutkind. 2010. Semaphorin 3E initiates antiangiogenic signaling through plexin D1 by regulating Arf6 and R-Ras. Mol. Cell. Biol. 30:3086–3098. doi:10.1128/MCB.01652-09.

Schindelin, J., I. Arganda-Carreras, E. Frise, V. Kaynig, M. Longair, T. Pietzsch, S. Preibisch, C. Rueden, S. Saalfeld, B. Schmid, J.-Y. Tinevez, D.J. White, V. Hartenstein, K. Eliceiri, P. Tomancak, and A. Cardona. 2012. Fiji: an open-source platform for biological-image analysis. Nat. Methods. 9:676–682. doi:10.1038/nmeth.2019.

Schneider, C.A., W.S. Rasband, and K.W. Eliceiri. 2012. NIH Image to ImageJ: 25 years of image analysis. Nat. Methods. 9:671–675.

Schweitzer, J.K., A.E. Sedgwick, and C. D’Souza-Schorey. 2011. ARF6-mediated endocytic recycling impacts cell movement, cell division and lipid homeostasis. Semin. Cell Dev. Biol. 22:39–47. doi:10.1016/j.semcdb.2010.09.002.

Scott, P.M., P.S. Bilodeau, O. Zhdankina, S.C. Winistorfer, M.J. Hauglund, M.M. Allaman, W.R. Kearney, A.D. Robertson, A.L. Boman, and R.C. Piper. 2004. GGA proteins bind ubiquitin to facilitate sorting at the *trans*-Golgi network. Nat. Cell Biol. 6:252–259. doi:10.1038/ncb1107.

Shevchenko, A., M. Wilm, O. Vorm, and M. Mann. 1996. Mass spectrometric sequencing of proteins silver-stained polyacrylamide gels. Anal. Chem. 68:850–858.

Shiba, Y., Y. Katoh, T. Shiba, K. Yoshino, H. Takatsu, H. Kobayashi, H.-W. Shin, S. Wakatsuki, and K. Nakayama. 2004. GAT (GGA and Tom1) domain responsible for ubiquitin binding and ubiquitination. J. Biol. Chem. 279:7105–7111. doi:10.1074/jbc.M311702200.

Steinberg, F., K.J. Heesom, M.D. Bass, and P.J. Cullen. 2012. SNX17 protects integrins from degradation by sorting between lysosomal and recycling pathways. J. Cell Biol. 197:219–230. doi:10.1083/jcb.201111121.

Takatsu, H., Y. Katoh, Y. Shiba, and K. Nakayama. 2001. Golgi-localizing, gamma-adaptin ear homology domain, ADP-ribosylation factor-binding (GGA) proteins interact with acidic dileucine sequences within the cytoplasmic domains of sorting receptors through their Vps27p/Hrs/STAM (VHS) domains. J. Biol. Chem. 276:28541–28545. doi:10.1074/jbc.C100218200.

Tang, Z., C. Li, B. Kang, G. Gao, C. Li, and Z. Zhang. 2017. GEPIA: a web server for cancer and normal gene expression profiling and interactive analyses. Nucleic Acids Res. 45:W98–W102. doi:10.1093/nar/gkx247.

Théry, M. 2010. Micropatterning as a tool to decipher cell morphogenesis and functions. J. Cell Sci. 123:4201–4213. doi:10.1242/jcs.075150.

Uemura, T., S. Kametaka, and S. Waguri. 2018. GGA2 interacts with EGFR cytoplasmic domain to stabilize the receptor expression and promote cell growth. Sci. Rep. 8:1368. doi:10.1038/s41598-018-19542-4.

Wang, S., C. Hu, F. Wu, and S. He. 2017. Rab25 GTPase: Functional roles in cancer. Oncotarget. 8:64591–64599. doi:10.18632/oncotarget.19571.

White, R.M., A. Sessa, C. Burke, T. Bowman, J. LeBlanc, C. Ceol, C. Bourque, M. Dovey, W. Goessling, C.E. Burns, and L.I. Zon. 2008. Transparent adult zebrafish as a tool for in vivo transplantation analysis. Cell Stem Cell. 2:183–189. doi:10.1016/j.stem.2007.11.002.

Winograd-Katz, S.E., R. Fässler, B. Geiger, and K.R. Legate. 2014. The integrin adhesome: from genes and proteins to human disease. Nat. Rev. Mol. Cell Biol. 15:273–288. doi:10.1038/nrm3769.

Ye, F., A.K. Snider, and M.H. Ginsberg. 2014. Talin and kindlin: the one-two punch in integrin activation. Front. Med. 8:6–16. doi:10.1007/s11684-014-0317-3.

Zhao, Y., and J.H. Keen. 2008. Gyrating clathrin: highly dynamic clathrin structures involved in rapid receptor recycling. Traffic. 9:2253–2264. doi:10.1111/j.1600-0854.2008.00819.x.

Zhu, Y., B. Doray, A. Poussu, V.P. Lehto, and S. Kornfeld. 2001. Binding of GGA2 to the lysosomal enzyme sorting motif of the mannose 6-phosphate receptor. Science. 292:1716–1718. doi:10.1126/science.1060896.

